# An *in silico* Cardiomyocyte Reveals the Impact of Changes in CAMKII Signalling on Cardiomyocyte Contraction Kinetics in Hypertrophic Cardiomyopathy

**DOI:** 10.1101/2023.05.11.540337

**Authors:** Ismail Adeniran, Hafsa Wadee, Hans Degens

## Abstract

Hypertrophic cardiomyopathy (HCM) is characterised by asymmetric left ventricular hypertrophy, ventricular arrhythmias and cardiomyocyte dysfunction that may cause sudden death. HCM is associated with mutations in sarcomeric proteins and is usually transmitted as an autosomal-dominant trait. The aim of this *in silico* study was to assess the mechanisms that underlie the altered electrophysiological activity, contractility, regulation of energy metabolism and crossbridge cycling in HCM at the single cell level. To investigate this, we developed a human ventricular cardiomyocyte model that incorporates electrophysiology, metabolism and force generation. The model was validated by its ability to reproduce the experimentally observed kinetic properties of human HCM induced by a) remodelling of several ion channels and Ca^2+^-handling proteins arising from altered Ca^2+^/calmodulin kinase II signalling pathways; and b) increased Ca^2+^ sensitivity of the myofilament proteins. Our simulation showed a decreased phosphocreatine to ATP ratio (−9%) suggesting a negative mismatch between energy expenditure and supply. Using a spatial myofilament half sarcomere model, we also compared the fraction of detached, weakly bound and strongly bound crossbridges in the control and HCM conditions. Our simulations showed that HCM has more crossbridges in force producing states than in the control condition. In conclusion, our model reveals that impaired crossbridge kinetics is accompanied by a negative mismatch between the ATP supply : demand ratio. This suggests that improving this ratio may reduce the incidence of sudden death in HCM.

## 1. Introduction

Hypertrophic cardiomyopathy (HCM) is the most common inherited cardiac disorder, affecting 1 in 500 individuals, and is clinically characterised by left ventricular (LV) hypertrophy, a decrease in the LV chamber size, heart failure, arrhythmias and sudden cardiac death at any age (Pasipoularides, 2018; Semsarian, 2018; Spudich, 2014). While the Ejection Fraction (EF) is preserved, the rate of relaxation is diminished (Haland et al., 2017; Marian and Braunwald, 2017; Spudich, 2014). It usually results from mutations of sarcomeric proteins; the sarcomere being the fundamental contractile unit of the cardiomyocyte (Pasipoularides, 2018; Spudich, 2014). Histopathological features include disorganized myocyte architecture, including disarray of myocyte fibres, intertwined hypertrophied myocytes, and focal or widespread interstitial fibrosis (Rowin and Fifer, 2021; Tejado and Jou, 2018).

Over the past two decades, it has been suggested that an energy deficit plays an essential role in the aetiology of HCM (Ormerod et al., 2016; Pasipoularides, 2018; Ranjbarvaziri et al., 2021; Wijnker et al., 2019). Most mutations in HCM lead to an increase in both the total force production and adenosine triphosphate (ATP) utilisation at the cellular level, consequently leading to a greater ATP demand by the cardiomyocytes (Ashrafian et al., 2003; Ranjbarvaziri et al., 2021). Notwithstanding this increased demand for ATP, both murine models (Lucas et al., 2003; Tardiff et al., 1999) and patients (Unno et al., 2009) with HCM exhibit an impaired, rather than enhanced, mitochondrial function and morphology. Despite significant advances in elucidating sarcomeric structure-function relationships and evidence linking impaired energy metabolism with the aetiology of HCM, there is still much to be discovered about the mechanisms that link altered cardiac energetics to HCM phenotypes due to insufficient cell and animal models and more importantly, human HCM patient studies (Li et al., 2018; Ranjbarvaziri et al., 2021).

Studies indicate that the pathogenesis of HCM involves a broad range of mechanisms; the primary mechanism of which is sarcomeric protein mutation. Molecular changes that occur in response to the changes in the sarcomere protein structure and function form a secondary mechanism, with histological and pathological phenotypes as a consequence of the molecular perturbations being tertiary mechanisms (Marian and Braunwald, 2017; Pasipoularides, 2018; Ranjbarvaziri et al., 2021; Rowin and Fifer, 2021). One such tertiary mechanism may be the complex remodelling of Ca^2+^/calmodulin-dependent protein kinase II (CAMKII)-dependent signalling (Coppini et al., 2018, 2013).

Advancements in understanding HCM will be facilitated by comprehensive models that can integrate complex factors influencing cardiomyocyte function. Indeed, previous models have made valuable contributions to understanding electromechanical characteristics in HCM. Computational modelling of cardiomyocytes in HCM revealed that the electromechanical features in HCM include delayed after-depolarisations, prolonged action potentials, Ca^2+^ overload, and hyperdynamic contraction, related to an elevation in Ca^2+^-related current densities and differences in Na^+^, K^+^ and Cl^-^-related current densities compared to normal conditions (Liu et al., 2013). The model of Pioner et al. (2023) found that electrophysiological changes (prolonged action potentials and Ca^2+^ transients) were responsible for a preserved twitch duration in a HCM-linked MYBPC3 mutation. Another model study saw in cardiomyocytes from HCM patients several abnormalities, including prolonged action potentials due to increased late sodium (I_NaL_) and calcium (I_CaL_) currents along with decreased repolarizing potassium currents, increased arrhythmias, abnormal calcium handling and increased CaMKII activity and phosphorylation (Coppini et al., 2013). Our *in silico* study applies a human ventricular cardiomyocyte model to assess how functional changes in CAMKII-dependent signalling impact electrophysiological activity, contractility and energy metabolism regulation at the level of the single cardiomyocyte in human HCM.

Our single cell model offers several advantages for studying HCM. First, it integrates electrophysiology, metabolism and force generation, providing a holistic understanding of the altered cellular dynamics in HCM. Second, our model incorporates experimentally observed kinetic properties of HCM involving ion channel remodelling and altered Ca^2+^ handling. This ensures that our simulations closely resemble the actual conditions in HCM. Third, the spatial myofilament half sarcomere model allows us to explore the distribution of crossbridges in force-producing states, providing insights into the mechanical aspects of HCM at a mesoscopic level. The model includes explicit representations of actin and myosin filaments. Although it is not an atomic scale model, the model seeks to represent spatial interactions between protein complexes that are thought to produce characteristic cardiac muscle responses at larger scales. Our work thus advances the HCM research by offering an encompassing approach to electrophysiology, metabolism, and force generation within a single-cell context.

## 2. Materials and Methods

### 2.1 Electromechanical, Mitochondrial Energetics Single Cell Model

For electrophysiology (EP), we employed the O’Hara-Rudy (ORd) human ventricular single cell model (O’Hara et al., 2011). We selected the ORd model because it is based on experimental data from human hearts and validated using cellular behaviours and mechanisms from an extensive dataset including measurements from over 100 undiseased human hearts. It accurately reproduces the electrical and membrane channel characteristics of human ventricular cells, as well as the transmural heterogeneity of ventricular action potentials (AP) across the ventricular wall. Furthermore, the ORd model replicates modulation of the rate of Ca^2+^ cycling by CaMK, which is particularly important in the context of this study. It also reproduces the Ca^2+^ *vs.* voltage-dependent inactivation of the L-type Ca^2+^ current.

To simulate cardiomyocyte contractile properties, we utilised the myofilament model developed by Tran et al. (Tran et al., 2010), an extension of the established Rice et al. cross-bridge cycling model of cardiac muscle contraction (Rice et al., 2008). Not only can this myofilament model replicate a variety of experimental data, such as steady-state force-sarcomere length (F-SL), force-Ca^2+^, and sarcomere length-Ca^2+^ relationships, but it can also reproduce many of the effects on force development due to MgATP, MgADP, Pi and H^+^ (Rice et al., 2008; Tran et al., 2010). In addition to these advantages, in the context of this study, the Tran et al. (Tran et al., 2010) model allowed us to assess the Hill coefficient to reproduce the change in the F-Ca^2+^ relationship in HCM seen by Coppini et al. (Coppini et al., 2013, 2018).

#### 2.1.1 Coupling the Electrophysiology and the Myofilament Models

The electrophysiology (EP) model was coupled with the myofilament model to produce an electromechanical cell model. The intracellular calcium concentration [𝐶𝑎^2+^]_𝑖_ from the EP model served as the link between the EP model and the myofilament model. Specifically, the dynamic [𝐶𝑎^2+^]_𝑖_ generated by the EP model in response to an AP was used as input to the myofilament model, which then calculated the quantity of Ca^2+^ bound to troponin.

The myoplasmic [𝐶𝑎^2+^]_𝑖_ concentration in the human ventricular myocyte electromechanical cell model is formulated as follows:

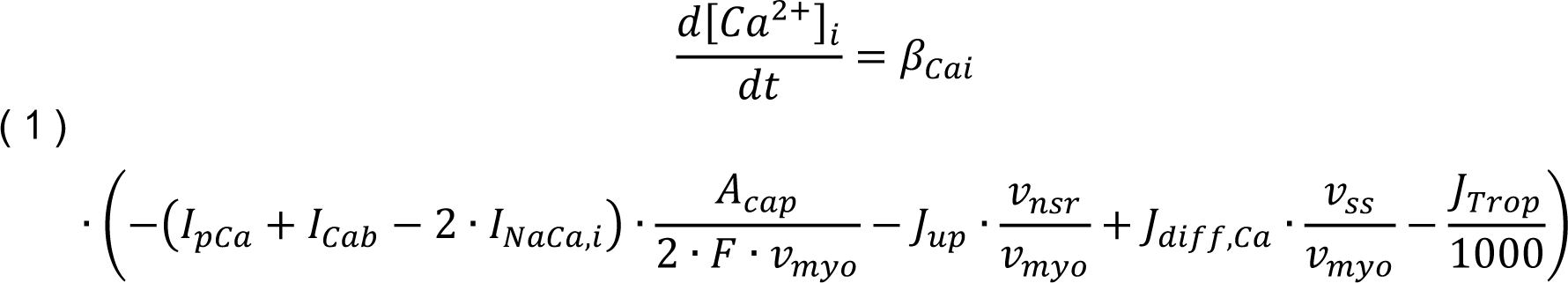

where *β_Cai_* is the buffer factor for [𝐶𝑎^2+^]_𝑖_, *I_pCa_* (μA/μF) is the sarcolemmal Ca^2+^ pump current, *I_Cab_* (μA/μF) is the Ca^2+^ background current, *I_NaCa,i_* (μA/μF) is the myoplasmic component of Na^+^/Ca^2+^ exchange current, *A_cap_* (cm^2^) is capacitive area, *F* (Coulomb/mol) is the Faraday constant, *v_myo_* (μL) is the volume of the myoplasmic compartment, *v_nsr_* (μL) is the volume of the sarcoplasmic reticulum compartment, *v_ss_* (μL) is the volume of the subspace compartment, *J_up_* (mM/ms) is the total Ca^2+^ uptake flux via the SERCA pump from myoplasm to the sarcoplasmic reticulum, *J_diff,Ca_* (mM/ms) is the diffusion flux of Ca^2+^ from the subspace to the myoplasm and *J_Trop_* (μM/ms) is the flux of *Ca^2+^* binding to troponin.

*β_Cai_* is formulated as:

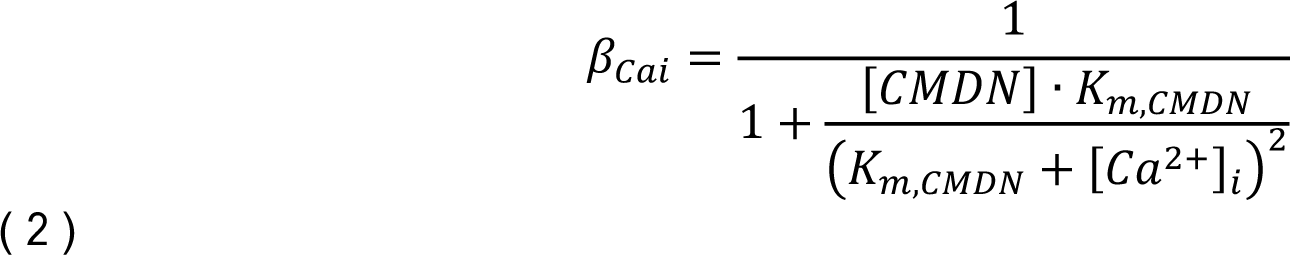

where [CMDN] is the calmodulin Ca^2+^ buffer in the myoplasm and *K_m,CMDN_* is the half-saturation concentration of calmodulin.

#### 2.1.2 Coupling the Electromechanical Model to the Mitochondrial Model

For cardiac mitochondrial bioenergetics, we then coupled this electromechanical single cell model to the thermokinetic, mitochondrial model of Cortassa et al. (Cortassa et al., 2003). This model describes the tricarboxylic acid (TCA) cycle, oxidative phosphorylation and mitochondrial Ca^2+^ handling. It can reproduce, qualitatively and semi-quantitatively, experimental data concerning mitochondrial bioenergetics, Ca^2+^ dynamics and respiratory control.

To consider mitochondrial energetics, modifications were made to the sarcolemmal Ca^2+^ pump current (*I_pCa_*) and the SERCA pump uptake flux (*J_up_*) of the electromechanical single cell model as follows:

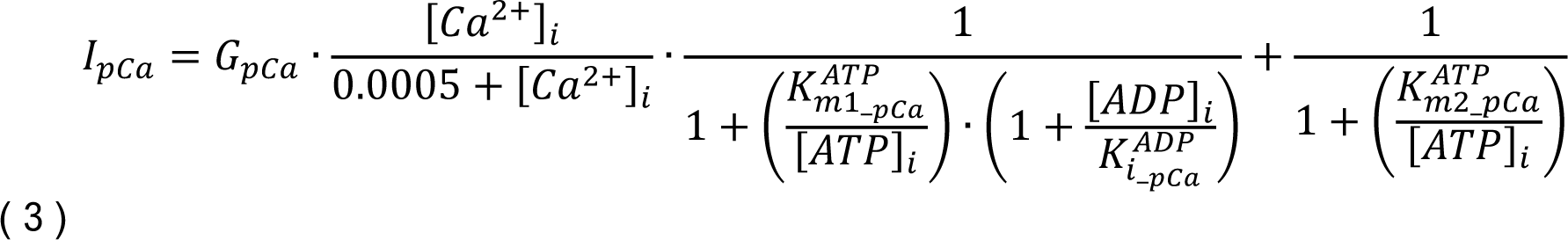

where *G_pCa_* is the maximum conductance of 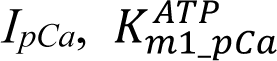 is the first ATP (adenosine triphosphate) half saturation constant for 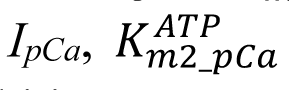 is the second ATP half saturation constant for 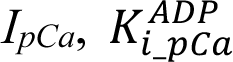 is the ADP inhibition constant for *I_pCa_*, *[ATP]_I_* is the excitation-contraction (EC) coupling linked ATP concentration and *[ADP]_I_* is the excitation-contraction (EC) coupling-linked ADP (adenosine diphosphate) concentration.

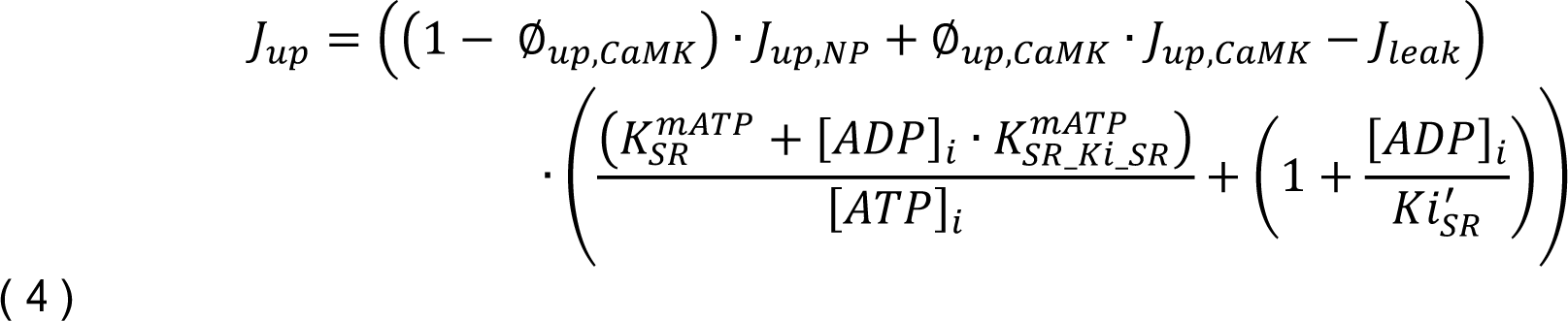

where *Ø_up,CaMK_* is the fraction of channels phosphorylated by CaMK, *J_up,NP_* is the non-phosphorylated Ca^2+^ uptake by CaMK via the SERCA pump, *J_up,CaMK_* is the CaMK phosphorylated Ca^2+^ uptake via SERCA pump and *J_leak_* is the leakage flux.

In addition to the above, the main driver of the interaction between the electromechanical model and the mitochondrial model is the [𝐶𝑎^2+^]_𝑖_ from the electrophysiology model. In the mitochondrial model, it influences processes such as ATP production. The mitochondrial model then feeds back to the electromechanical model to influence Ca^2+^ cycling and Ca^2+^ handling.

### 2.2 HCM Electromechanical Single Cell Model

To develop the human electromechanical single cell model of a HCM cardiomyocyte, we modified the EP and myofilament models to reflect the experimental data from Coppini et al. (2013, 2018) on the electromechanical profile of cardiomyocytes from 26 HCM patients. We made the following changes to the electromechanical single cell model (see Table 1).

**Table 1:**
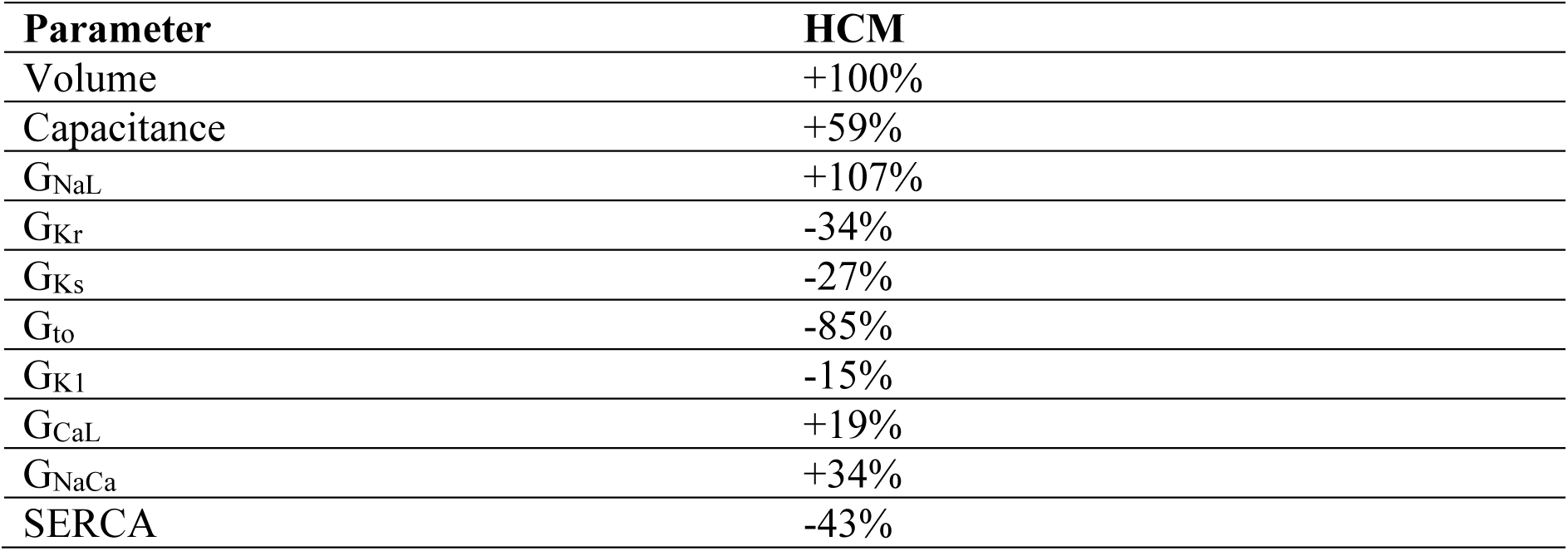
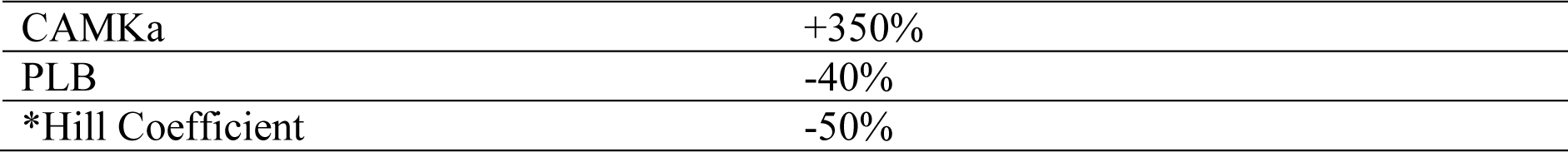
Hypertrophic cardiomyopathy (HCM) functional changes (Coppini et al., 2013, 2018) made to the electromechanical single cell model. *Change made to the myofilament model of the coupled single model. GNaL is the maximum conductance of the late sodium current, GKr is the maximum conductance of the rapid delayed rectifier K^+^ current, GKs is the maximum conductance of the slow delayed rectifier K^+^ current, GKr is the maximum conductance of the rapid delayed rectifier K^+^ current, Gto is the maximum conductance of the transient outward K^+^ current, GK1 is the maximum conductance of the inward rectifier K^+^ current, GCaL is the maximum conductance of the Ca^2+^ current through the L-type Ca^2+^ channel, GNaCa is the maximum conductance of the Na^+^/K^+^ exchange current, SERCA is the sarco/endoplasmic reticulum Ca^2+^-ATPase pump, CaMKa is the fraction of active Ca^2+^/calmodulin-dependent protein kinase II binding sites, PLB is phospholamban and the Hill coefficient is a measure of the Ca^2+^ sensitivity of the cardiomyocyte.

### 2.3 Spatially Explicit, Half-sarcomere Multifilament Model

To further investigate pathological behaviour in HCM, we also employed a spatially explicit, multifilament computational model of the half sarcomere (Powers et al., 2018; Williams et al., 2010, 2012, 2013). This model is comprised of eight actin filaments and four myosin filaments arranged in a three-dimensional double-hexagonal lattice (see Figures 1a and 1b). It simulates an infinite lattice using periodic (toroidal) boundary conditions (see Figure 1b). Each cross-bridge is modelled as a two-spring system consisting of a torsional and linear component. Linear springs are connected in series between each node of cross-bridge crowns for the myosin and actin filaments (see Figure 1c). Each myosin filament contains 60 crowns of three myosin motors, resulting in 180 cross-bridges per myosin filament.

**Figure 1:**
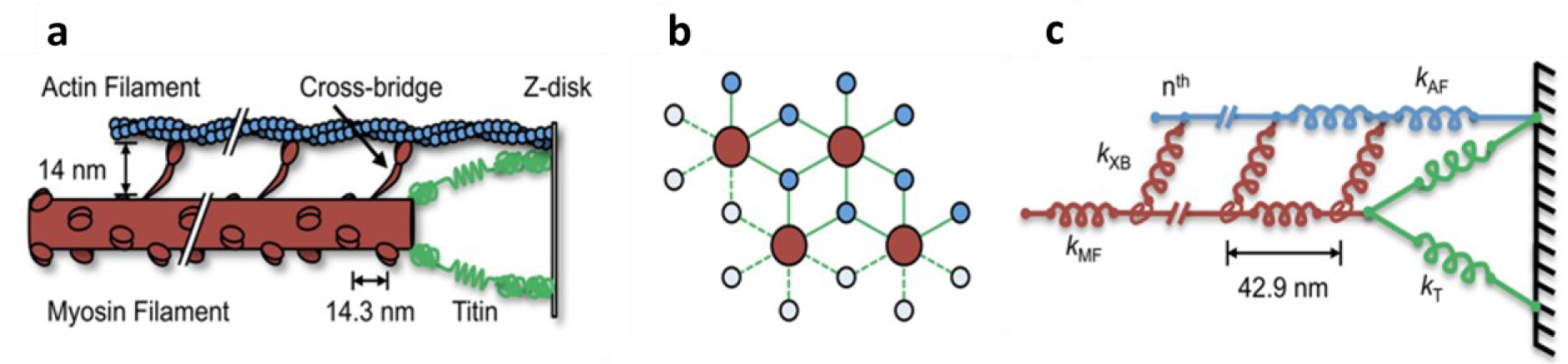
Spatially explicit, half sarcomere model (modified from (Powers et al., 2018)). (a) Schematic of the half sarcomere showing myosin filament (red), actin filament (blue), titin (green), Z-disk (black) and crossbridges (red). (b) Spatial hexagonal arrangement of myosin and actin filaments. (c) Mathematically, the half sarcomere is modelled as an array of springs: myosin and crossbridges (red), actin (blue) and titin (green).

Importantly, the model possesses two noteworthy characteristics that distinguish it from other existing models. First, it integrates torsional springs and lever-arm mechanisms that expose a correlation between step size and lattice spacing. Second, this lever-arm mechanism generates both radial and axial forces of similar magnitude, mirroring experimental findings (Williams et al., 2010).

As input to the multifilament, half sarcomere model, we digitised [𝐶𝑎^2+^]_𝑖_ transients from control and HCM cardiomyocytes recorded in two separate studies. The first set comes from 26 HCM patients in (Coppini et al., 2013, 2018) (see Figure 2a). The second set is from 93 HCM patients with a myosin binding protein C (MYBPC3) mutation (Pioner et al., 2023). Mutations to MYBPC3 are the most common cause of HCM (Helms et al., 2020).

**Figure 2:**
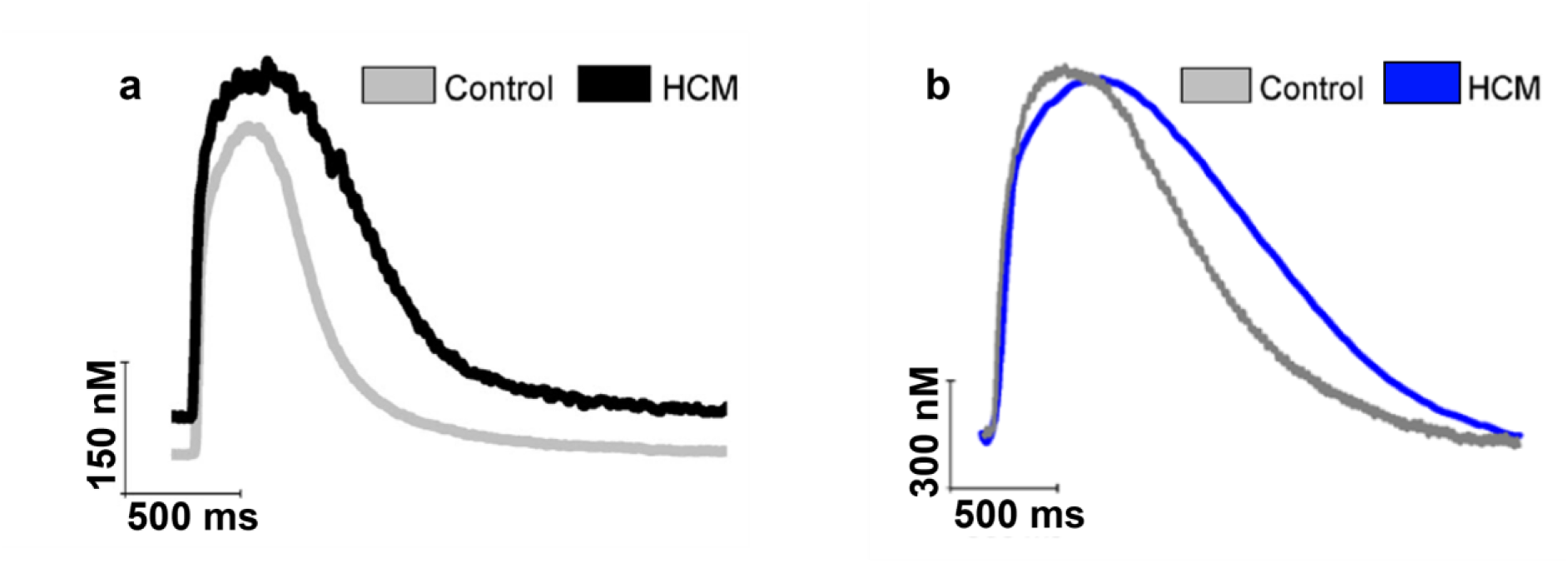
[𝐶𝑎^2+^]_𝑖_ transients recorded from control and HCM cardiomyocytes input into the multifilament half-sarcomere model. (a) Transients recorded from in current clamp mode in a study on 26 HCM patients (Coppini et al., 2013). (b) Transients recorded from study on 96 patients with a MYBPC3 mutation (Pioner et al., 2023).

In each case, the maximum [𝐶𝑎^2+^]_𝑖_ for the control condition was set equivalent to an actin permissiveness of 1 during crossbridge cycling while the minimum [𝐶𝑎^2+^]_𝑖_ for the control condition was set equivalent to an actin permissiveness of 0.2. Values of actin permissiveness were proportionally adjusted between these values during the time course simulation of the pre-recorded profiles for both control and HCM conditions. Each time point of each calcium transient profile was run for 100 ms until reaching steady state, at which point the steady values of the active force and the fraction of crossbridges in the strongly bound, weakly bound and detached states were recorded (see Figure 5).

### 2.4 Parameter Values, Initial conditions and Numerical Integration Methods

All the parameter values and initial conditions for the EP model, the myofilament model, the mitochondrial energetics model and spatial half sarcomere model are as described in the corresponding manuscripts and their supplementary materials: (O’Hara et al., 2011) for the electrophysiology model, (Tran et al., 2010) for the myofilament model, (Cortassa et al., 2003) for the mitochondrial model and (Powers et al., 2018) for the half sarcomere model. The changes made to the ORd EP model as well as the 50% reduction to the Hill coefficient in the myofilament model are detailed in Table 1. These changes were made as they align with the experimentally observed changes between control and HCM patients (Coppini et al., 2018, 2013). Numerical integration of the ordinary differential equations (ODEs) was performed using the LSODA ODE differential equation solver (Hindmasrsh et al., 2005).

## 3. Results and Discussion

### 3.1 Model Validation: HCM Electromechanical Single Cell

To validate the HCM single cell model, we computed a steady-state force-Ca^2+^ (F-pCa) relationship for a sarcomere length (SL) of 2.2 μm and compared the outcome with experimental data (Coppini et al., 2013). The results are shown in Figure 3. The model reproduced the differences in total and passive forces between control and HCM, which matched experimental data (inset).

**Figure 3:**
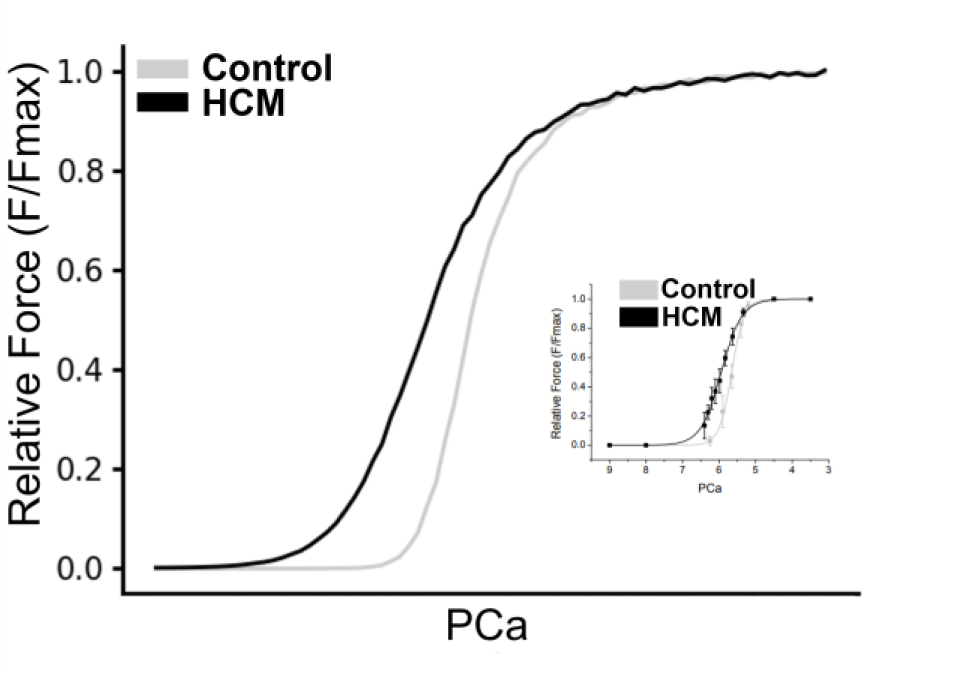
Force-pCa Relationship. Simulated Force-pCa relationship in control and HCM. Relative force is normalised to maximum value that can be attained. Inset: Experimental force-pCa relation of skinned HCM and control preparations obtained from human patients (Coppini et al., 2013).

### 3.2 Functional Electrophysiological Consequences of HCM

Figure 4 shows the electromechanical consequences of HCM in a free-running cell from the epicardium. HCM increased the action potential duration (APD_90_) from 231 ms to 342 ms (see Figures 4a, 5a and 6a). This is qualitatively similar to what was experimentally observed in HCM cardiomyocytes (Coppini et al., 2013).

**Figure 4:**
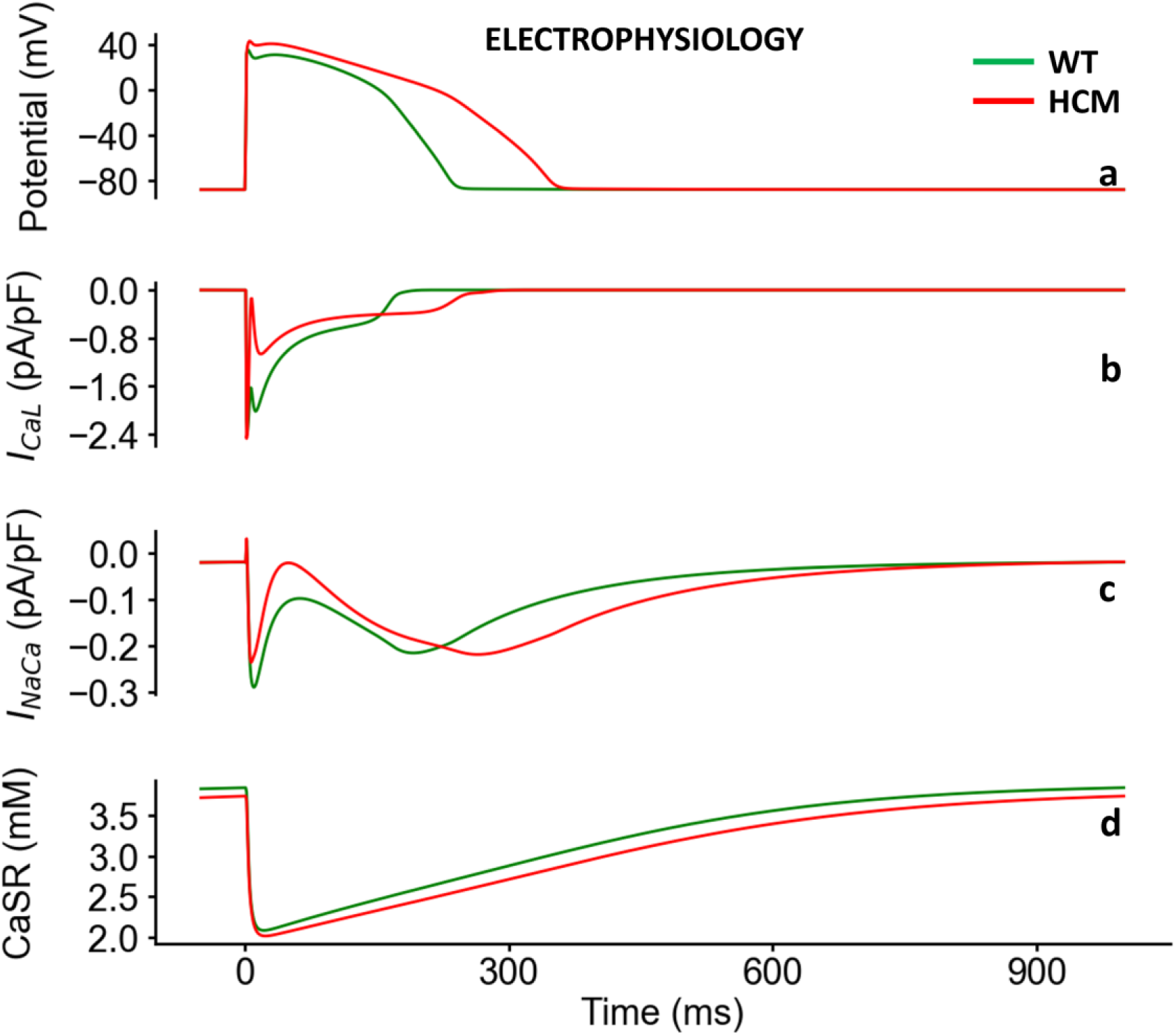
Electrophysiological properties under control (green) and HCM (red) at 1 Hz cycle length. **(a):** Epicardial action potential in HCM and control. **(b)** L-type Ca^2+^ channel current (ICaL). **(c)** Sodium/calcium exchange current (INaCa). **(d)** Ca^2+^ release and re-uptake from and to the sarcoplasmic reticulum.

**Figure 5:**
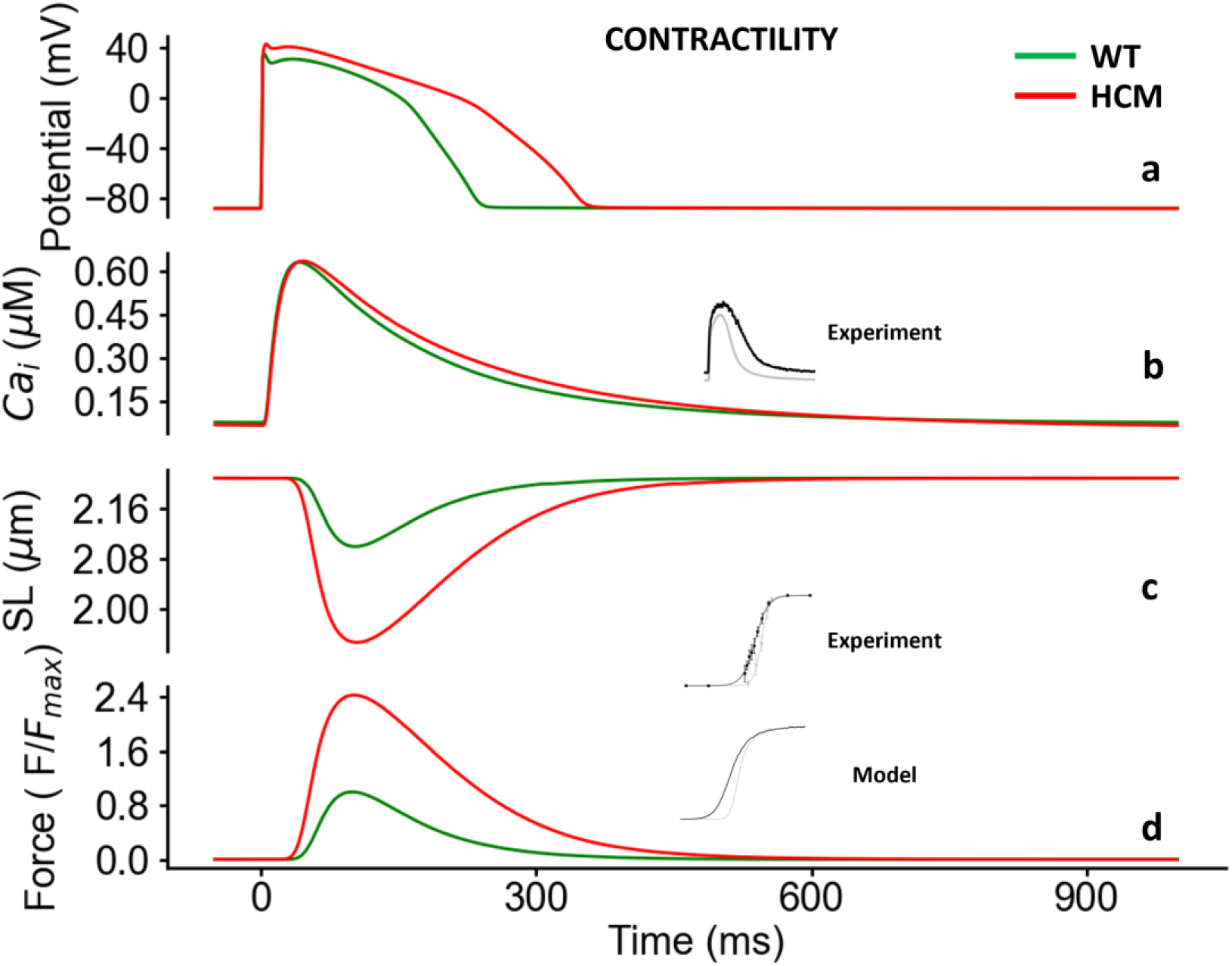
Contractile properties under control (green) and HCM (red) at 1 Hz cycle length. **(a):** Epicardial action potentials in HCM and control. **(b)** [𝐶𝑎^2+^]_𝑖_ transients. **(c)** Sarcomere length (SL). **(h)** Active force. Values are normalised to Control maximum active force.

**Figure 6:**
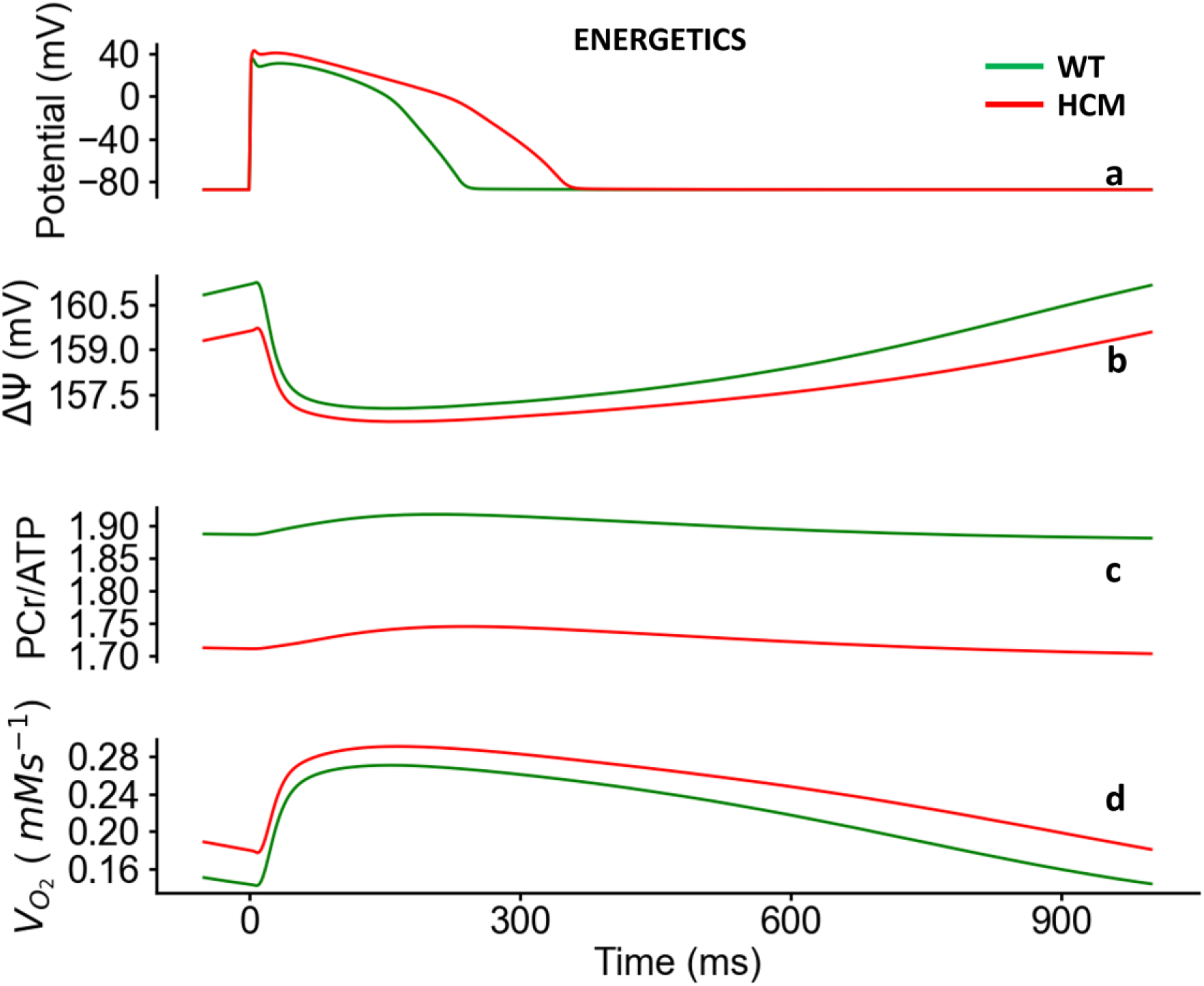
Bioenergetical properties under control (green) and HCM (red) at 1 Hz cycle length. **(a):** Epicardial action potentials in HCM and control. **(b)** Time course of change of mitochondrial membrane potential. **(c)** Phosphocreatine to ATP (PCr/ATP ratio. **(d)** Oxygen consumption rate (V02).

The L-type Ca^2+^ current density (l_CaL_) increased in HCM at the onset of AP depolarisation but reduced in amplitude during the plateau phase (see Figure 4b). Although the intracellular Ca^2+^ concentration [𝐶𝑎^2+^]_𝑖_ amplitude was similar in HCM and control (see Figure 5b), its kinetics were slower in HCM. The slowed kinetics of [𝐶𝑎^2+^]_𝑖_ (see Figure 5b) coupled with the transient increase in l_CaL_, led to an increased amplitude of the reverse-mode of the Na^+^/Ca^2+^ exchanger (NCX) while leaving its forward mode unchanged (see Figure 4c). The initial concentration of in the Sarcoplasmic Reticulum (SR) is lower in HCM but in absolute terms, the release is the same as in WT (see Figure 4d). The SR Ca^2+^ content is less in HCM. These factors, along with the decreased conductances of repolarising K^+^ currents (see Table 1; G_Kr_, G_Ks_, G_to_, G_K1_), contributed to the prolongation of the action potential in HCM (see Figure 4a, 5a, 6a).

With every heartbeat, the [𝐶𝑎^2+^]_𝑖_ undergoes an order of magnitude change, while the resulting force may change by three or more orders of magnitude. This biological design is believed to be motivated by the need for significant changes in developed force to efficiently pump blood.

When incomplete relaxation occurs, or delayed relaxation (see Figure 5d), a residual force, and hence pressure in the ventricle, inhibits proper filling during diastole, resulting in the ejection of a smaller amount of blood during systole. In contrast, cardiac cells produce relatively small changes in [𝐶𝑎^2+^]_𝑖_ because ATP is required to actively decrease [𝐶𝑎^2+^]_𝑖_ with each heartbeat. The steep nonlinearity of the F-pCa relationship illustrates the small change in [𝐶𝑎^2+^]_𝑖_ and the large changes in force. A steep nonlinearity enables a proportionally larger change in force for a much smaller relative change in [𝐶𝑎^2+^]_𝑖_ per heartbeat (Chung et al., 2016; Sun and Irving, 2010). Figure 3 shows that the force-pCa relationship is left-shifted in HCM patients indicating increased Ca^2+^ sensitivity, which resulted in greater sarcomere length shortening (see Figure 5c) and greater developed force in HCM with a lesser magnitude of [𝐶𝑎^2+^]_𝑖_ (see Figure 5d).

The mitochondrion is a major organelle storing Ca^2+^ and regulates the [𝐶𝑎^2+^]_𝑖_ homeostasis with the SR (Santo-Domingo and Demaurex, 2010). The potential energy derived from the mitochondrial membrane potential (ΔΨ) is essential for the generation of ATP during oxidative phosphorylation (Santo-Domingo and Demaurex, 2010; Zorova et al., 2018). ΔΨ was reduced in HCM (see Figure 6b), implying less potential energy available for the generation of ATP by the enzyme ATP-synthase, resulting in a potential mismatch between energy (ATP) demand and supply. This is indicated by a decreased phosphocreatine to ATP ratio (−9%) (see Figure 6c) and an increased oxygen consumption rate, or respiration rate (see Figure 6d). The ratio of phosphocreatine/ATP (PCr/ATP) reflects the availability of phosphocreatine and is a measure of the energy state of the myocardium (Del Franco et al., 2022; Henry et al., 2022; Neubauer et al., 1997). The impaired cardiac energetics in HCM seen in our model is consistent with a multiomics profile study of 27 HCM patients and 13 controls (Ranjbarvaziri et al., 2021), providing validation of our developed human, electromechanical, energetic single cell model.

### 3.3 Sensitivity Analysis of the HCM Electromechanical Model Parameters

Table 1 shows the parameters that were modified in the control electromechanical model to obtain the HCM electromechanical model. Given the complexity of the integrated models, we performed a sensitivity analysis to identify the key parameters that have the most significant impact on the results. We therefore systematically modified each parameter in the control electromechanical model at a time to determine its effects. The results are shown and discussed below.

#### 3.3.1 Effect of the Changes on Membrane Potential

The only parameter that had a significant effect on the action potential duration was the 34% reduction of I_Kr_ (see Figure 7). This caused the APD_90_ to increase from 231 ms to 328 ms, which is still shorter than the full HCM electromechanical model’s APD_90_ of 342 ms. I_Kr_ is a repolarising K^+^ current that is active during the plateau phase of the action potential, and its reduction results in a delayed repolarisation or a prolonged APD.

**Figure 7:**
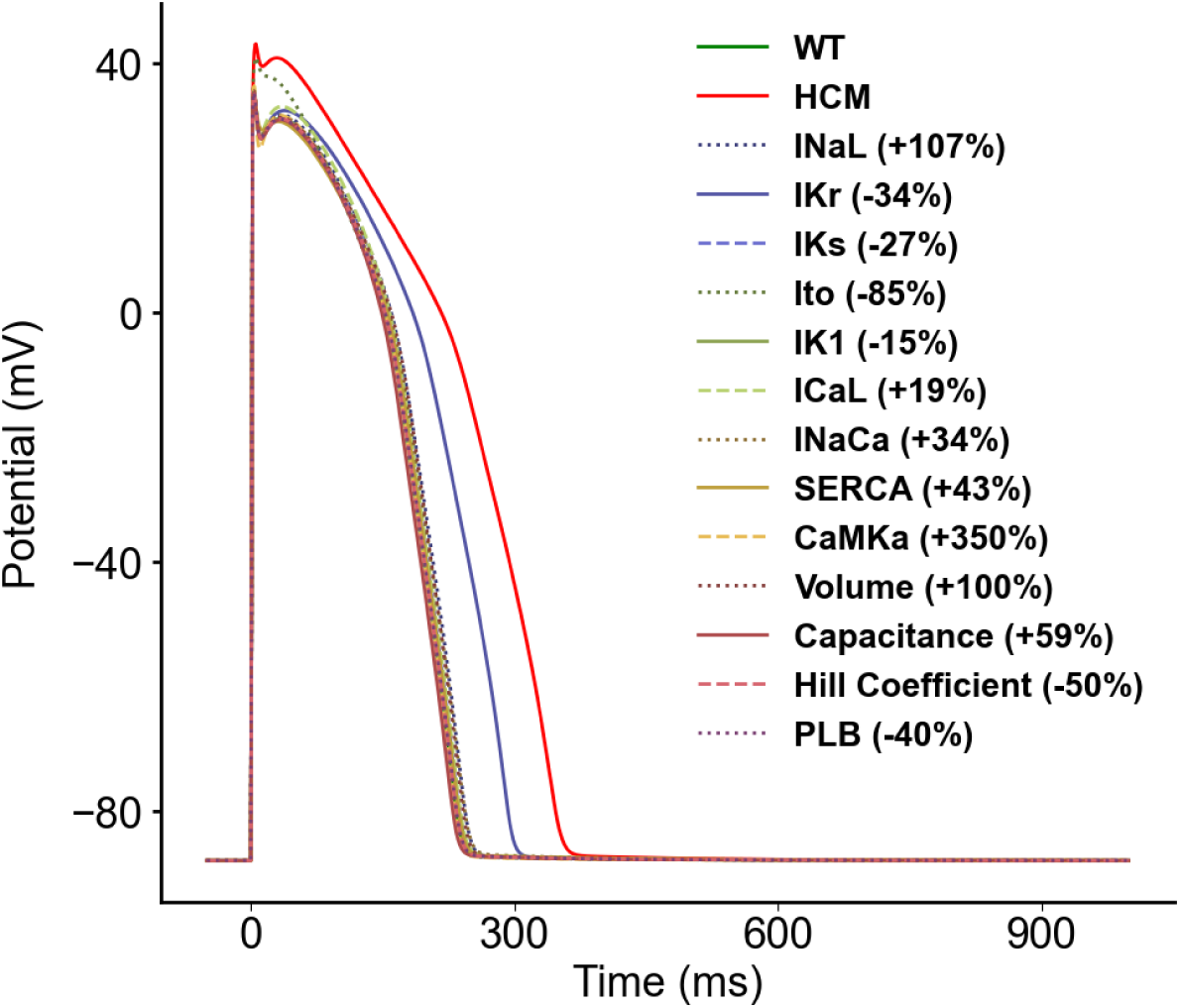
Sensitivity analysis of the model parameters compared to control and HCM at 1Hz cycle length on the membrane potential.

#### 3.3.2 Effect of the Changes on CaSR

In WT, the initial [Ca^2+^] in the SR was ∼3.7 mM and had a maximal reduction to ∼2.1 mM during the action potential (see Figure 8). In HCM, the initial [Ca^2+^] of the SR was ∼3.6 mM with a maximal reduction to ∼2 mM during the action potential (see Figure 8). All parameter changes had an effect approximately between those of WT and HCM, i.e., a maximal Ca^2+^ release of ∼50% except for changes in CaMK, and SERCa. CaMK had the greatest effect (see Figure 8). While the change in CaMK also had a maximal release of ∼50%, the maximal [Ca^2+^] was elevated to ∼5.2 mM.

**Figure 8:**
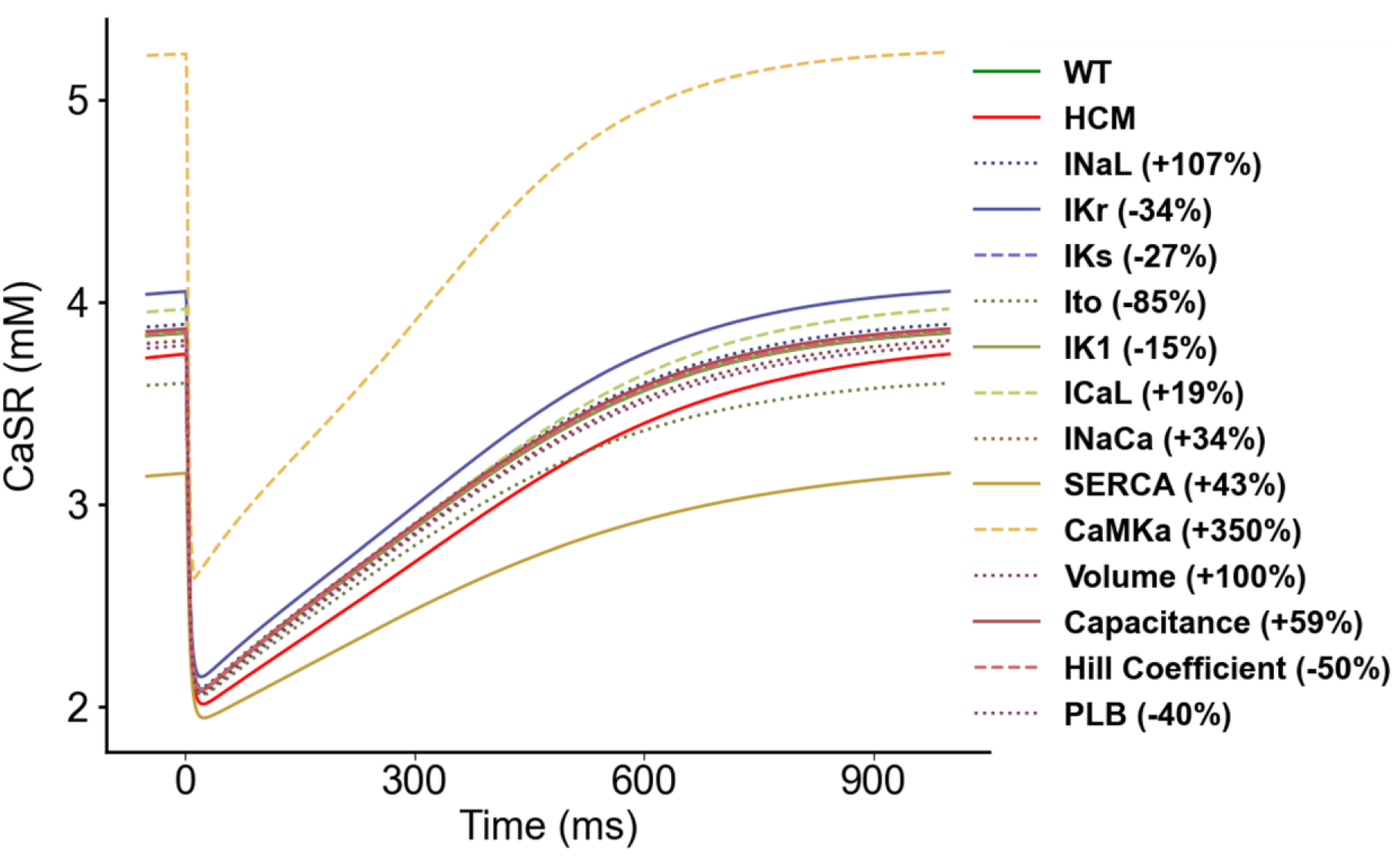
Sensitivity analysis of the model parameters compared to control and HCM at 1Hz cycle length on ICaL.

In HCM, CaMK is elevated by 350%. This elevation would be expected to result in an increased SR [Ca^2+^] by stimulating uptake of Ca^2+^ by SERCA into the SR, leading to a higher SR Ca^2+^ load. The higher SR Ca^2+^ content then provides more releasable Ca^2+^ with each subsequent action potential. This release is further enhanced by promotion of Ca^2+^ release by direct action of CaMK on the ryanodine receptor (RyR) and activation of CaMK by the calcium-calmodulin complex. Therefore, the higher cytosolic Ca^2+^ levels not only activate CaMK but also create a positive feedback loop. While this enhances the heart’s pumping ability, it introduces a potential risk of Ca^2+^-driven arrhythmias due to the heightened release of Ca^2+^ and increased cytosolic Ca^2+^ concentrations.

#### 3.3.3 Effect of the Changes on [𝐶𝑎^2+^]_𝑖_

The increase in CaMK had the greatest effect on the [𝐶𝑎^2+^]_𝑖_, increasing the amplitude from ∼0.62 µM to ∼0.9 µM (see Figure 9). Since CaMK phosphorylates and activates RyR, the channels open more easily resulting in greater SR Ca^2+^ release that will increase the amplitude of [𝐶𝑎^2+^]_𝑖_. Additionally, CaMK phosphorylates and inhibits phospholamban, an inhibitor of the SERCA pump. This disinhibition of the SERCA pump allows greater SR calcium reuptake by SERCA accelerating the [𝐶𝑎^2+^]_𝑖_ decay (see Figure 9) as Ca^2+^ is pumped back into the SR. The increased SR Ca^2+^ reuptake also replenishes the SR Ca^2+^ stores maintaining a higher SR Ca^2+^ content. This further augments the amount of releasable Ca^2+^ (see Figure 8).

**Figure 9:**
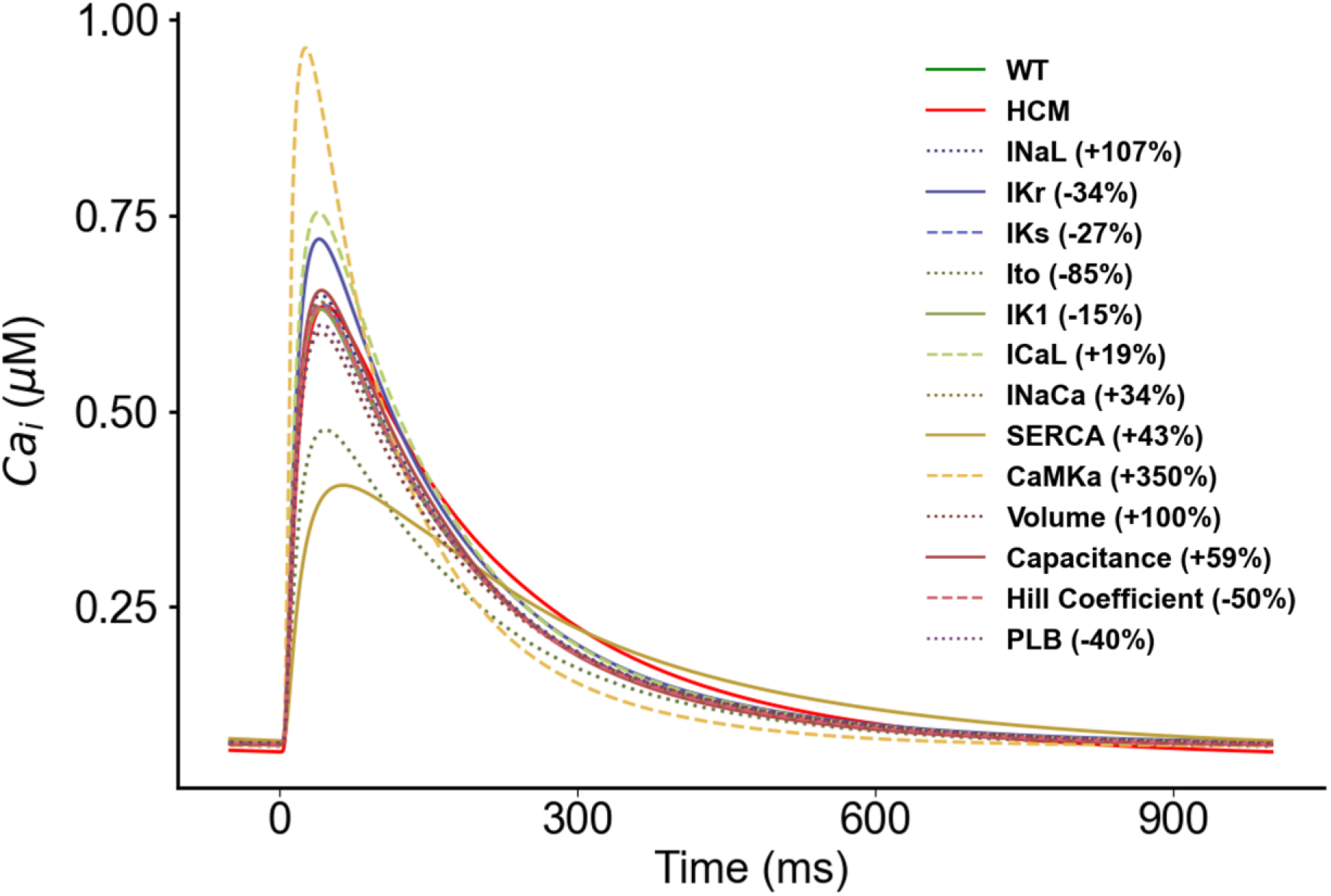
Sensitivity analysis of the model parameters compared to control and HCM at 1Hz cycle length on [𝐶𝑎^2+^]_𝑖_.

I_CaL_ also increased the [𝐶𝑎^2+^]_𝑖_ amplitude from ∼0.62 µM to ∼0.75 µM. I_CaL_ provides the influx of trigger Ca^2+^ that initiates calcium-induced calcium release from the SR. Therefore, an increase in I_CaL_ would result in greater Ca^2+^ influx during the AP plateau, contributing to a higher [𝐶𝑎^2+^]_𝑖_ level. The increased trigger Ca^2+^ activates more RyR channels leading to greater SR Ca^2+^ release and an increase in the amplitude of [𝐶𝑎^2+^]_𝑖_ (see Figure 9).

The 34% increase in I_NaCa_ reduced the [𝐶𝑎^2+^]_𝑖_ amplitude from ∼0.62 µM to ∼0.45 µM (see Figure 9). I_NaCa_ is responsible for the extrusion of Ca^2+^ from the cardiomyocyte. Therefore, an increase in I_NaCa_ activity means that more Ca^2+^ is pumped out of the cardiomyocyte reducing the [𝐶𝑎^2+^]_𝑖_ (see Figure 9).

Most of the other parameter changes had an effect approximately intermediate between WT and HCM (see Figure 9).

#### 3.3.4 Effect of the Changes on the Active Force

The 350% increase of CaMK increased the active force of WT 2.5-fold (see Figure 10). CaMK phosphorylates and activates the RyR Ca^2+^ release channels, which results in greater Ca^2+^ release from the SR. The increased SR Ca^2+^ release amplifies [𝐶𝑎^2+^]_𝑖_, providing more Ca^2+^ to bind to troponin C and initiate myofilament crossbridge cycling. CaMK phosphorylates and inhibits phospholamban, which disinhibits SERCA. This increases SR Ca^2+^ reuptake. Faster SR Ca^2+^ reuptake allows quicker relaxation (see Figure 10) and replenishes SR Ca^2+^ stores, further increasing releasable calcium (see Figure 8). Overall, this leads to an increase in the active tension and force generation by cardiomyocytes (see Figure 10).

**Figure 10:**
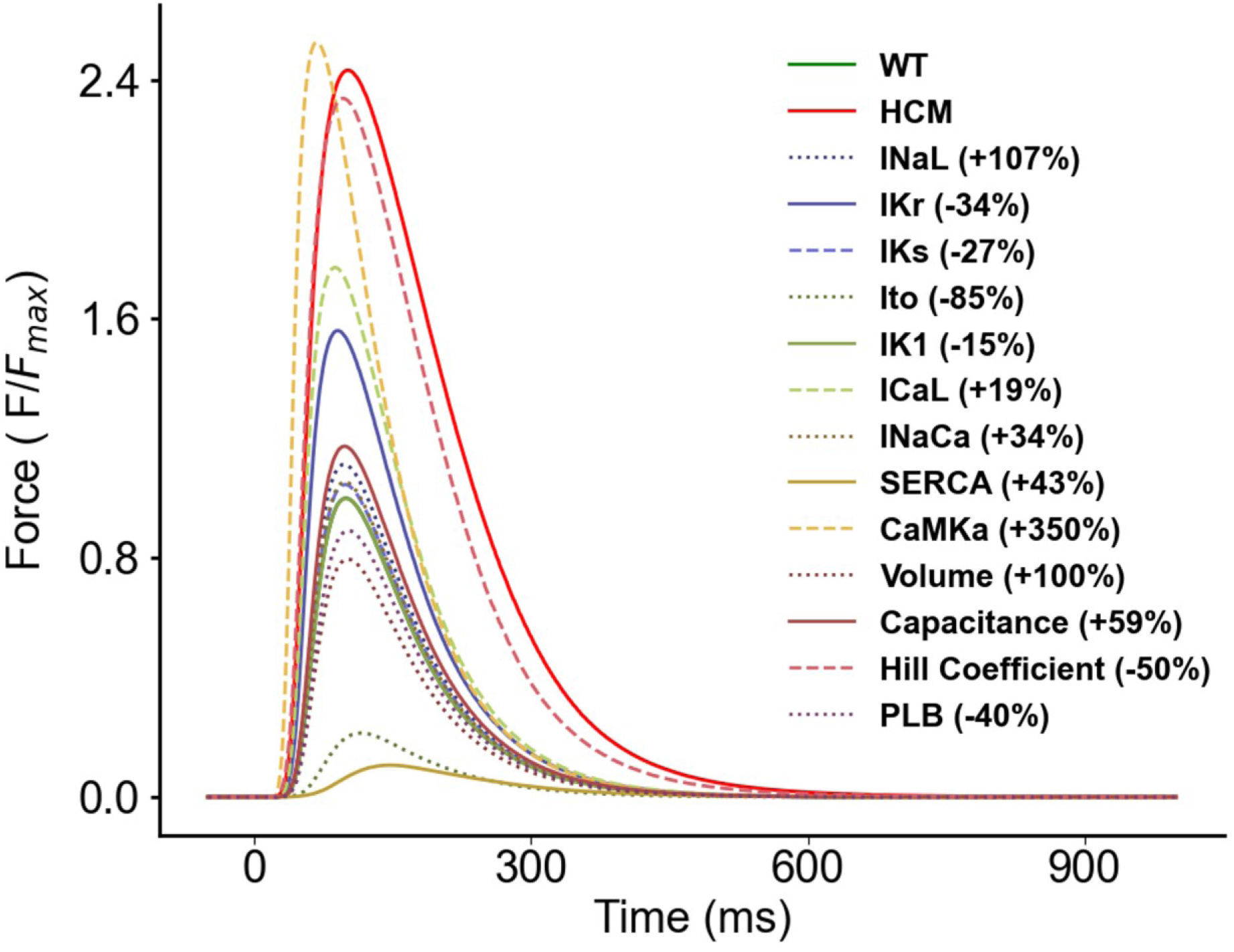
Sensitivity analysis of the model parameters compared to control and HCM at 1Hz cycle length on active force. IK1 is superimposed on WT.

The 50% decrease in the Hill coefficient also increased the active force of WT ∼2.3-fold (see Figure 10). The Hill coefficient quantifies the steepness of the Force-Ca^2+^ relationship and quantifies the cooperative binding of Ca^2+^ to troponin C which activates the myofilaments in cardiomyocytes. With reduced steepness and consequently, lower cooperativity, binding of Ca^2+^ to troponin C occurs more gradually. Without cooperative effects, Ca^2+^ dissociates more gradually from troponin C thereby slowing relaxation and increasing active force (see Figure 10).

The 19% increase in I_CaL_ increased the active force of WT ∼1.8-fold (see Figure 10). I_CaL_ provides Ca^2+^ influx during the AP plateau, acting as the trigger for calcium-induced calcium release from the SR. Increased I_CaL_ density would result in increased SR Ca^2+^ release. This would lead to an increase in active tension and force generation (see Figure 10). The higher Ca^2+^ level would also hasten relaxation (see Figure 10) by providing more substrate for removal, mainly by the sodium-calcium exchanger (NCX).

I_NaCa_ reduced the active force to 10% of WT (see Figure 10). It primarily functions by exchanging three Na^+^ into the cell for one Ca^2+^ out of the cell. An increase in I_NaCa_ activity (43% under HCM) leads to high Ca^2+^ extrusion, reducing the [𝐶𝑎^2+^]_𝑖_. By lowering [𝐶𝑎^2+^]_𝑖_, the increase in I_NaCa_ indirectly influences contractility. Reduced [𝐶𝑎^2+^]_𝑖_ during diastole means that there will be less Ca^2+^ available for the next contraction, leading to a decrease in contractile force (see Figure 10).

I_Kr_ is reduced by 34% in HCM. This reduction slows repolarisation and prolongs the APD, which lengthens the period of Ca^2+^ influx via the action of I_CaL_. The increased Ca^2+^ influx can enhance calcium-induced calcium release from the SR and augment [𝐶𝑎^2+^]_𝑖_. The higher [𝐶𝑎^2+^]_𝑖_ leads to increased Ca^2+^ binding to troponin C and greater myofilament crossbridge formation. This results in an increase in peak active tension and force generation and slower relaxation (see Figure 10).

The other parameter changes had moderate effect on active force compared to HCM when all the changes were incorporated.

#### 3.3.5 Effect of the Changes on the Mitochondrial Membrane Potential (ΔΨ)

All the parameters had effects that were approximately intermediate between WT and HCM except for CaMK, SERCA and I_to_ (see Figure 9). CaMK is increased by 350% in HCM, SERCA by 43% while I_to_ is reduced by 85% (see Table 1).

The greatly elevated CaMK hyperphosphorylates and overactivates the RyR. This leads to excessive Ca^2+^ release from the SR and Ca^2+^ overload in the cytosol and mitochondria. Mitochondrial Ca^2+^ overload can cause opening of the mitochondrial permeability transition pore (mPTP). Opening of the mPTP dissipates the mitochondrial membrane potential (ΔΨ) (see Figure 11) and impairs ATP production. The reduction in ΔΨ prevents normal Ca^2+^ uptake into the mitochondria, further disrupting cellular Ca^2+^ regulation. CaMK also increases metabolism and ATP demand; combining this with reduced ATP production over time depletes energy stores.

**Figure 11:**
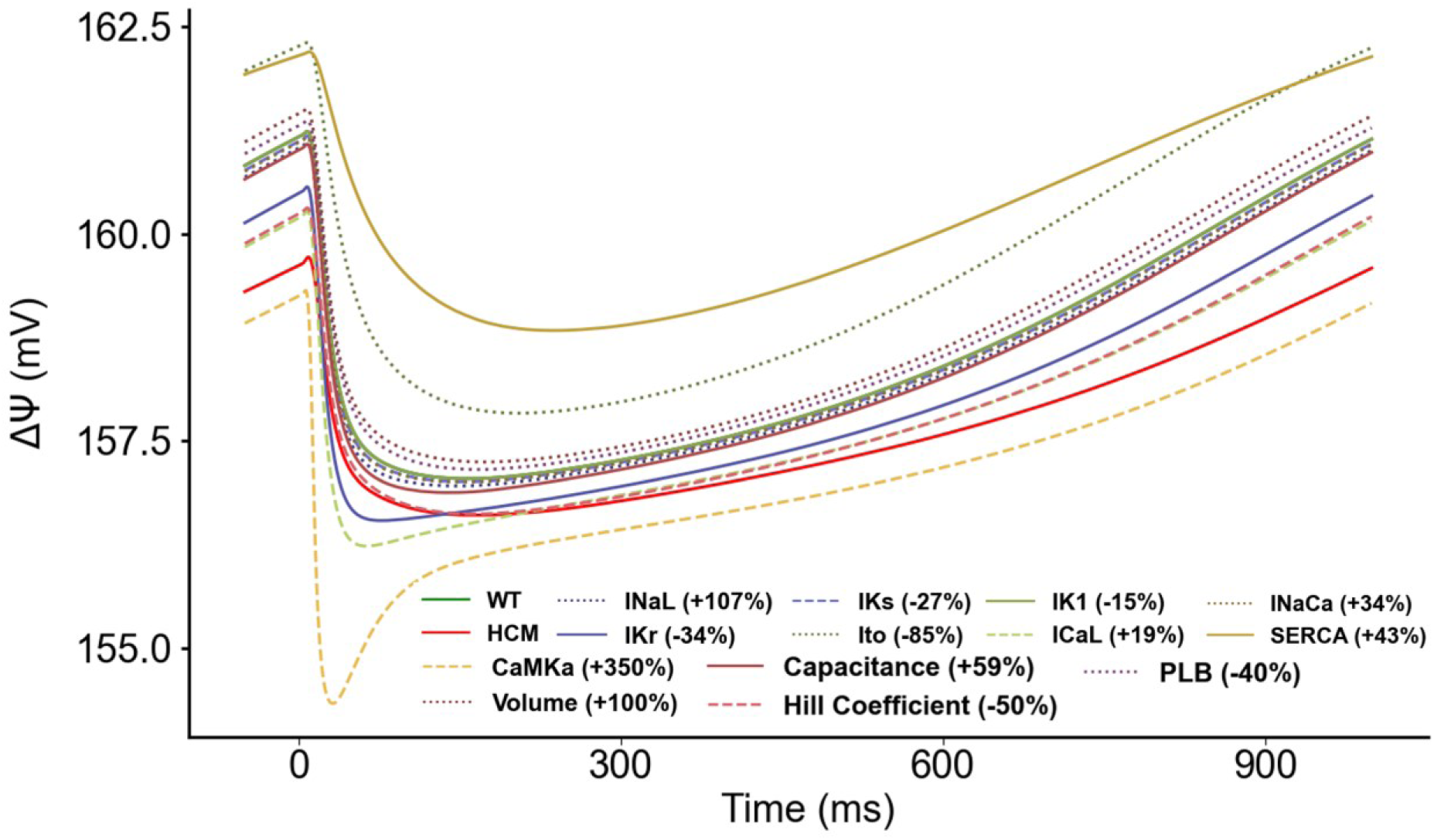
Sensitivity analysis of the model parameters compared to control and HCM at 1Hz cycle length on mitochondrial membrane potential. IK1 is superimposed on WT.

The 43% increase in SERCA had the opposite effect in that it augmented ΔΨ (see Figure 11). Increased SERCA activity accelerates Ca^2+^ reuptake into the SR during relaxation. Faster Ca^2+^ clearance from the cytosol prevents Ca^2+^ overload and reduces entry of Ca^2+^ into the mitochondria. Since excessive mitochondrial Ca^2+^ can trigger opening of the mPTP, the increased SERCA activity sustains ΔΨ by depolarising the inner mitochondrial membrane and preventing Ca^2+^ overload (see Figure 11). The 85% reduction of I_to_ in HCM also augmented ΔΨ . This was because loss of I_to_ prolonged the AP and increased Ca^2+^ influx primarily through I_CaL_ activity. This leads to Ca^2+^ overload in the cytosol and subsequently in the mitochondria. Mitochondrial Ca^2+^ overload triggers opening of the mPTP, and opening the mPTP dissipates ΔΨ and impairs ATP production.

#### 3.3.6 Effect of the Changes on the PCr/ATP Ratio

CaMK, SERCA and I_to_ had the most significant effect on the PCr/ATP ratio (see Figure 12). CaMK reduced the PCr/ATP ratio from ∼ 1.9 in WT to ∼1.5. SERCA increased the ratio from ∼1.9 to ∼ 2.2 while I_to_ increased it from ∼1.9 to ∼2.1. All the other parameters had effects that were intermediate between WT and HCM or close to WT (see Figure 12). CaMK is increased by 350% in HCM, SERCA by 43% while I_to_ is reduced by 85% (see Table 1).

**Figure 12:**
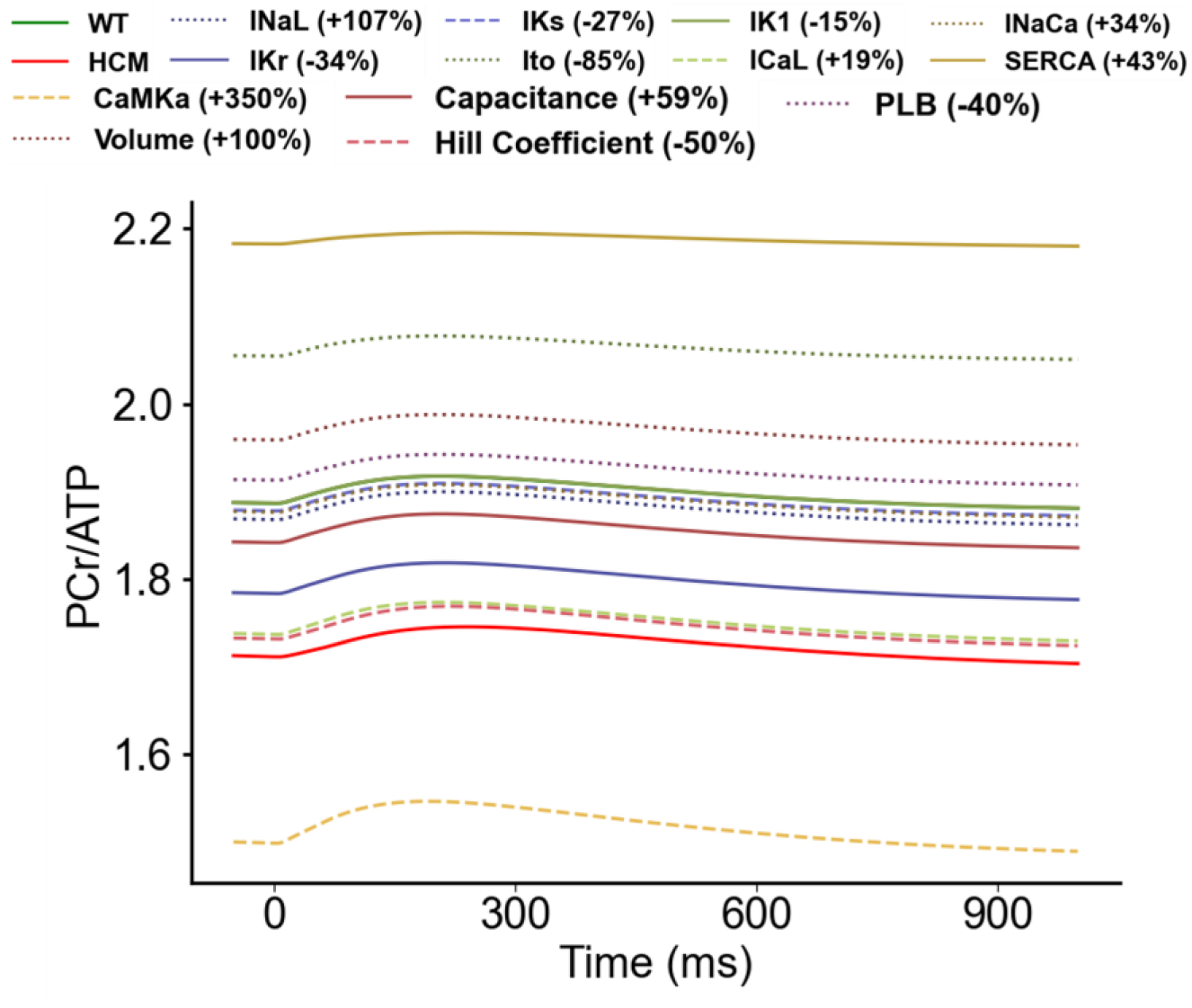
Sensitivity analysis of the model parameters compared to control and HCM at 1Hz cycle length on PCr/ATP ratio. IK1 is superimposed on WT.

As a possible mechanism for CaMK, the 350% increase in CaMK activity leads to excessive Ca^2+^ release from the SR, causing cytosolic and mitochondrial Ca^2+^ overload. This may lead to increased ATP demand to support enhanced contractile activity and ion transport. This higher ATP demand then leads to the utilisation of PCr stores to regenerate ATP and consequently a reduction in the PCR/ATP ratio (see Figure 12).

The 85% reduction in I_to_ leads to a prolongation of the APD, increasing Ca^2+^ influx through I_CaL_. This causes Ca^2+^ overload thereby activating SERCA, which consumes ATP. The resulting increase in ATP consumption leads to an increase in the PCr/ATP ratio. An additional mechanism is that excess mitochondrial Ca^2+^ triggers sustained opening of the mPTP, which depolarises the inner mitochondrial membrane and impairs ATP production.

The 43% increase in SERCA activity means that more Ca^2+^ is pumped back into the SR. This can reduce [𝐶𝑎^2+^]_𝑖_, implying that less energy is required to maintain ion gradients, such as NCX activity. With reduced energy expenditure on calcium handling, more ATP is available for regeneration of PCr. These effects can ultimately contribute to the increased PCr to ATP ratio (see Figure 12).

#### 3.3.7 Effect of the Changes on the Oxygen Consumption Rate (V_O2_)

CaMK, SERCA and I_to_ had the most significant effects on V_O2_ (see Figure 13). CaMK had a maximum peak value of 0.43 mM·s^-1^, I_to_ had a maximum amplitude of 0.24 mM·s^-1^ and SERCA’s maximum amplitude was 0.21 mM·s^-1^. The maximum amplitude for WT was 0.27 mM·s^-1^. The other parameters had maximum amplitudes that were comparable with WT or intermediate between WT and HCM (see Figure 13).

**Figure 13:**
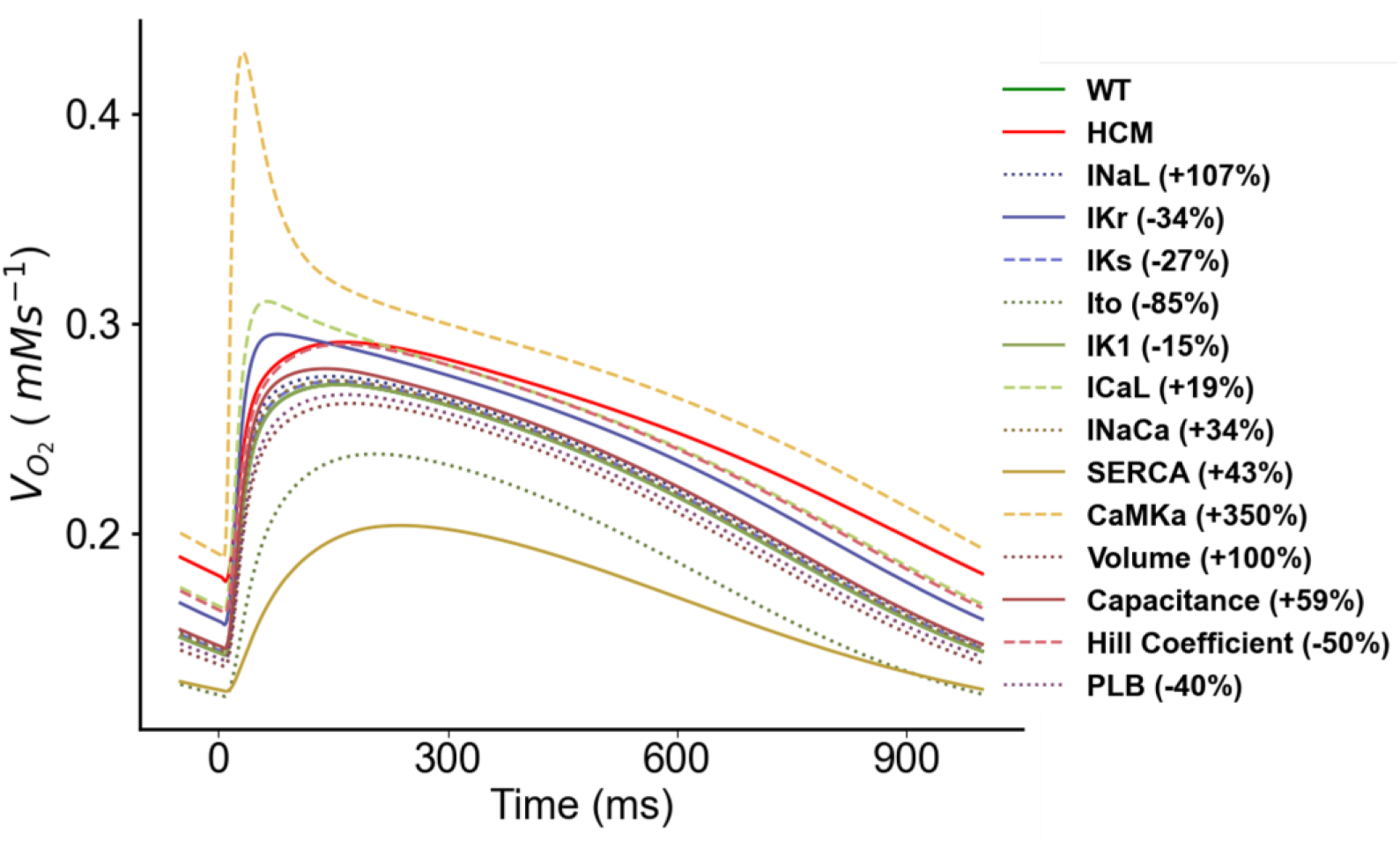
Sensitivity analysis of the model parameters compared to control and HCM at 1Hz cycle length on oxygen consumption rate (VO2). IK1 is superimposed on WT.

### 3.4 Functional Consequences of HCM on Crossbridge Cycling

Using the spatial, multifilament, half sarcomere model (Powers et al., 2018; Williams et al., 2010, 2012, 2013), we further investigated the functional consequences of HCM on crossbridge cycling. The half sarcomere model is a three-state model with states for crossbridge attachment, force generation and detachment. The transitions between these states determine both the forces experienced by each cross-bridge and the rate of ATP utilisation associated with the number of times each cross-bridge detaches from the thin filament and hydrolyses a subsequent ATP for subsequent binding and force generation. There is no thin filament regulation in the model as Ca^2+^ regulation is not considered. So, we could not couple it to an electrophysiological model as was done in the electromechanical model in sections 2.1 and 3.2. Instead, Ca^2+^ activation/regulation is considered indirectly in the model through actin permissiveness, which can be set to values between 0 (non-permissive) and 1 (fully permissive). This implies that, in this setting, troponin/tropomyosin interactions are combined into a single variable. While this simplification allows for meaningful qualitative observations, it may not fully capture the complexity of actual molecular mechanisms.

We used as input to the model the experimental [𝐶𝑎^2+^]_𝑖_ transient profiles of control and HCM patients (see Figure 2). These were digitised and each timepoint of the pertinent profile was run until steady state, after which the steady state value was recorded. This was done throughout the time course of each [𝐶𝑎^2+^]_𝑖_ transient profile. In each case, the minimum value of the [𝐶𝑎^2+^]_𝑖_ transient in the control condition was mapped to an actin permissiveness of 0.2 and the maximum [𝐶𝑎^2+^]_𝑖_ transient value was mapped to an actin permissiveness of 1. [𝐶𝑎^2+^]_𝑖_ transient values were mapped proportionally between these actin permissiveness values.

Figure 14a shows the active force in control and HCM for the [𝐶𝑎^2+^]_𝑖_ transient profile (inset). The force in both conditions follows closely the input [𝐶𝑎^2+^]_𝑖_ transient profile. This should not be the case, i.e., the force should not depend exactly on the [𝐶𝑎^2+^]_𝑖_ transient profile as experimental transients contain artifacts from a nonlinear response of the Ca^2+^-sensing dyes. The [𝐶𝑎^2+^]_𝑖_ transient and the force should be dynamically generated from a dynamic, free running cell (model) based on the interplay of ion channels and other cellular components.

Unfortunately, as discussed above, the half sarcomere model does not consider Ca^2+^ regulation directly but only through actin permissiveness. Thus, we were unable to couple the half sarcomere model to an electrophysiology model. This has certain implications that merit consideration. The lack of direct Ca^2+^ regulation may affect the dynamic interplay between ion channels and cellular components, which could affect how well our model mimics the complexities of a real cell. Additionally, we acknowledge that experimental [𝐶𝑎^2+^]_𝑖_ transient profiles inherently contain artifacts, and our model, by not directly incorporating Ca^2+^ regulation, may not fully capture the dynamic responses expected in a free-running cellular environment. Nevertheless, the results are qualitatively sound.

Figures 14b, 14c and 14d show that on average, in the rest state, the control condition has ∼97.5% of crossbridges in the detached state (HCM ∼96%), ∼1.5% in the weakly bound state (HCM ∼2.4%) and ∼1.5% in the strongly bound state (HCM ∼1.6%).

**Figure 14:**
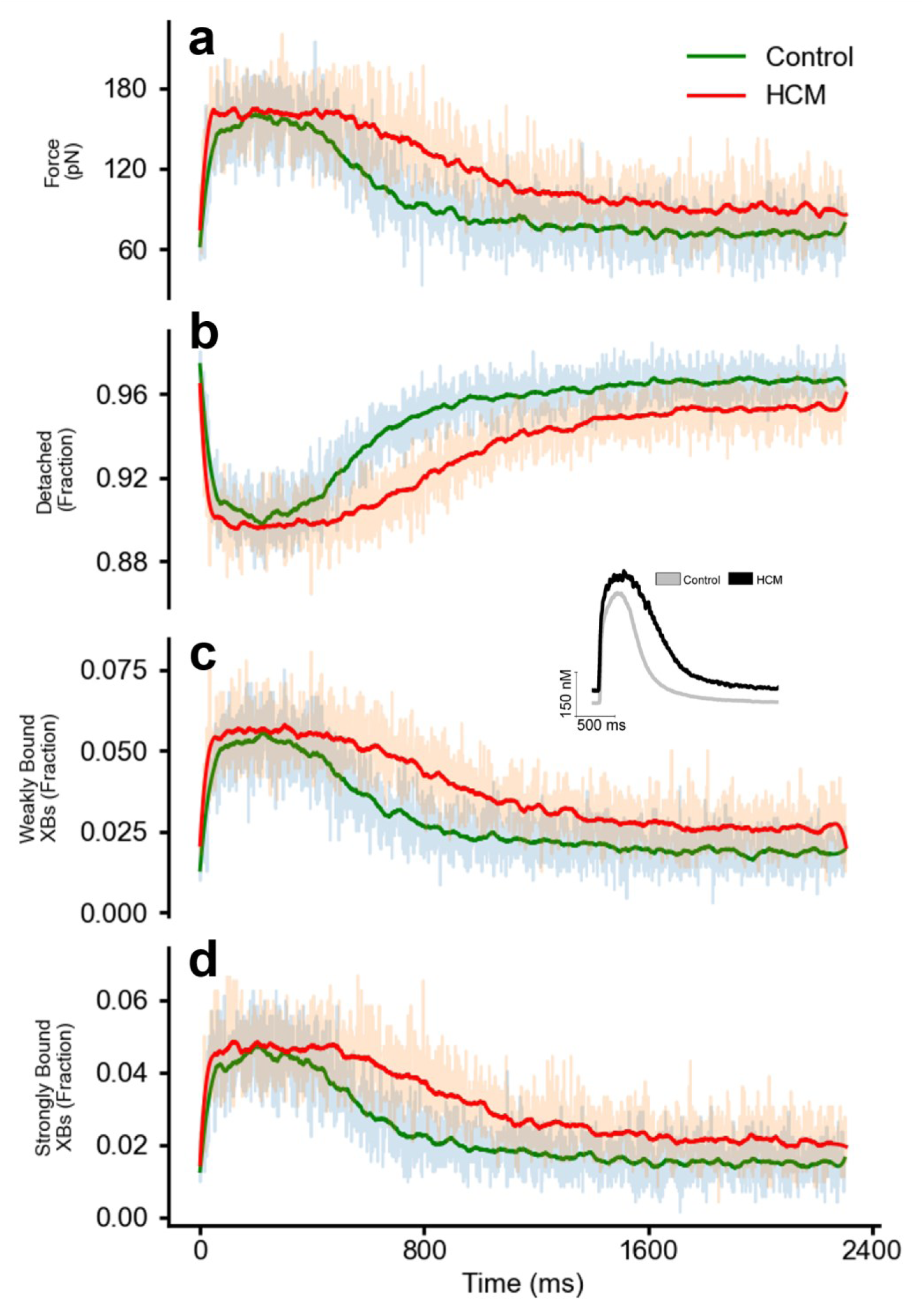
Crossbridge states in control and HCM with the spatial multifilament, half sarcomere model. (a) The developed force in control (green) and HCM (red). (b) The fraction of crossbridges in the detached state during the time course of the [𝐶𝑎^2+^]_𝑖_ transient profile. (c) The fraction of crossbridges in the weakly bound state during the time course of the [𝐶𝑎^2+^]_𝑖_ transient profile. (d) The fraction of crossbridges in the strongly bound state during the time course of the [𝐶𝑎^2+^]_𝑖_ transient profile.

In the weakly bound state, the myosin head has broken ATP into ADP (adenosine diphosphate) and phosphate (Pi). During this phase, the head has just attached to actin but has not yet released ADP or Pi. In this state, the myosin is only partially bound to actin and little or no force is produced. The myosin head then rotates into a strongly bound configuration, while ADP and Pi are released. In this state, myosin performs the primary force-producing “power stroke” of the cross-bridge cycle as it pulls on the actin filament (Lymn and Taylor, 1971). This implies that even at rest, HCM cardiomyocytes are more primed to generate force.

Throughout the time course of the active force profiles (see Figure 14a), the HCM condition has more crossbridges in the weakly bound and strongly bound states. At about 1000 ms, the control condition has ∼96% detached crossbridges, while this is 94% in HCM ∼94% (see Figure 14b). For the weakly bound it is ∼2.5% crossbridges for controls and 3.75% for HCM (see Figure 14c) and for the strongly bound ∼2% and ∼3%, respectively (see Figure 14d). This proportion is maintained throughout the rest of the time course of the profiles. This shows that from heartbeat to heartbeat, in the HCM condition more crossbridges are in the weakly bound (very low force generating) and in the strongly bound (force generating) states compared to the control condition.

Figure S1 in the supplementary material shows the result from a second [𝐶𝑎^2+^]_𝑖_ transient profile from HCM patients with a MYBPC3 mutation. It shows a similar trend to the results in Figure 14 and corroborates that HCM indeed has a higher proportion of crossbridges in the force generating states both during contraction and in the relaxed state, compared to control conditions.

### 3.5 Implications and Significance of Major findings

In this study, we developed a human ventricular electromechanical model that incorporated experimentally observed electrophysiological and mechanical changes during HCM. We also used a half sarcomere to investigate the effects of reported HCM Ca^2+^ transients to investigate the mechanisms underlying altered crossbridge cycling in HCM. Our simulations implicated CaMK, SERCA and I_to_ as key contributors to HCM pathophysiology. In HCM, CaMK is elevated by 350%, SERCA is increased by 43% and I_to_ is reduced by 85%. Identifying elevated CaMK, elevated SERCA and reduced I_to_ as major players in HCM pathophysiology provides crucial insights into the underlying molecular and cellular mechanisms of the disease. An CaMK increase by itself leads to excessive Ca^2+^ release from the SR and Ca^2+^ overload in the cytosol and mitochondria. While a compensatory elevated SERCA activity enhances Ca^2+^ re-uptake into the SR, it also provides more Ca^2+^ for SR release during subsequent beats, helping maintain contractility. However, the 85% decrease in I_to_ serves to prolong the APD and provides more time for Ca^2+^ influx into the cytosol. This continuous cycle is at the cost of greater energy demand beyond supply.

Consequently, strategies to reduce CaMK and SERCA, and increase I_to_ activity or modulate their downstream effects can potentially offer new treatment options for individuals with HCM. In addition, elevated CaMK levels in tandem with increased SERCA activity and reduced I_to_ levels could potentially help identify individuals with HCM at higher risk or with more severe disease and guide clinical management decisions.

### 3.6 Relevance to Previous Studies

Liu et al. (2023) developed a computational model to investigate the behaviour of electromechanical coupling in the human left ventricular cardiomyocyte and explore the roles of various factors in the pathological manifestations of HCM cardiomyocyte and their changes in response to drug treatment. Their study revealed that in addition to the characteristic changes in I_CaL_ and intracellular Na^+^ concentration, almost all other ion channel current densities and intracellular ion and molecule concentrations suffered from marked changes. This agrees with our model of HCM based on the experimental changes observed by Coppini et al. (Coppini et al., 2013, 2018) (see Table 1).

Pioner et al. (2023) used a computational model to test the mechanism responsible for a preserved twitch duration with the underlying prolonged AP and Ca2^+^ transients in a HCM-linked MYBPC3 mutation. They found that electrophysiological changes (prolonged action potentials and Ca^2+^ transients) appear to counterbalance the faster cross-bridge cycling, ultimately preserving the amplitude and duration of cardiac contraction, at the expense of cardiac electrical stability and diastolic function. This supports our result of the spatial half sarcomere model showing more crossbridges in the bound state in HCM (see Figure 14).

Forouzandehmehr et al. (2022) developed a metabolite-sensitive computational model of human induced pluripotent stem cell-derived cardiomyocytes (hiPSC-CMs) electromechanics to investigate the pathology of R403Q HCM mutation and the effect of Mavacamten (MAVA), Blebbistatin (BLEB), and Omecamtiv mecarbil (OM) on the cell mechano-energetics. Their model captured the prolonged contractile relaxation due to the R403Q mutation without assuming any further modifications such as an additional Ca^2+^ flux to the thin filaments. However, their model did not employ a spatial model and did not consider other electrophysiological changes to the HCM cardiomyocyte.

### 3.7 Limitations of These Simulations

The limitations of the ORd model (O’Hara et al., 2011), the Tran et al. model (Tran et al., 2010; Rice et al., 2008) and the mitochondrial model (Cortassa et al., 2003) have been discussed in the respective model papers. Our single cell simulations share these limitations.

In addition to these limitations, there is no thin filament regulation in the spatial myofilament model as Ca^2+^ regulation is not considered. Therefore, we could not couple it to an electrophysiological model as was done with the single cell electromechanical model in sections 2.1 and 3.2. Instead, Ca^2+^ activation/regulation is considered indirectly in the model through actin permissiveness, which can be set to values between 0 (non-permissive) and 1 (fully permissive). This implies that, in this setting, troponin/tropomyosin interactions are combined into a single variable.

While this simplification allows for meaningful qualitative observations, it may not fully capture the complexity of the actual cellular mechanisms. This has potential implications for the overall accuracy and biological significance of our model. The absence of direct Ca^2+^ regulation, a fundamental aspect of muscle contraction, may influence the dynamic interplay between ion channels and cellular components. This could impact how well the model mimics the intricacies of a living cell.

We also used as input to the spatial half sarcomere model pre-recorded [𝐶𝑎^2+^]_𝑖_ transients. Hence, the force output depends exactly on the [𝐶𝑎^2+^]_𝑖_ transient profile. This should not be the case as experimental transients contain artifacts from a nonlinear response of the Ca^2+^-sensing dyes. The [𝐶𝑎^2+^]_𝑖_ transient and the force should be dynamically generated from a dynamic, free running cell (model) based on the interplay of ion channels and other cellular components. Unfortunately, as discussed above, the half sarcomere model does not consider Ca^2+^ regulation directly but only through actin permissiveness. Thus, we were unable to couple the half sarcomere model to an electrophysiology model.

It may have also been beneficial to use a greater number of samples for the pre-recorded [𝐶𝑎^2+^]_𝑖_ transients but more samples which compared control with HCM were difficult to source. Ultimately, even if more samples were used, the results would still be subject to the problem alluded to in the above paragraph, i.e., the force output depending exactly on the [𝐶𝑎^2+^]_𝑖_ transient profile. Consequently, more targeted experiments and experimental data focused solely on this goal should be performed and collected.

At present, the modifications made to the EP and myofilament models to reflect the electromechanical profile of HCM cardiomyocytes are based on experimental data from Coppini et al. (2013, 2018). It would be beneficial to have a more comprehensive validation of the modified model against additional experimental data to ensure its accuracy in capturing HCM characteristics.

While it is important that potential limitations of the models used in this study are made explicit, these do not fundamentally affect the conclusions that can be drawn on likely mechanisms by which HCM alters electrophysiological activity, contractility, energy metabolism regulation and crossbridge cycling.

## Conclusions

HCM cardiomyocytes exhibited greater contractility (Figure 4) due to the combination of a) slowed [𝐶𝑎^2+^]_𝑖_ decay kinetics; b) reduced SERCA activity and phospholamban (PLB); c) greater myofilament Ca^2+^ sensitivity and d) a higher proportion of crossbridges in force-generating states compared to the control condition. All of these combined to produce, or were derived from, Ca^2+^ accumulation in the cytoplasm. This greater cardiomyocyte contractility is what enables the heart, and in particular the left ventricle, to maintain a relatively normal ejection fraction despite the wall hypertrophy in HCM. Our results show that a relatively normal function is maintained in HCM due to Ca^2+^ handling abnormalities but to achieve normal or close to normal function, the cardiomyocytes have a greater maximal oxygen consumption rate. Our results also implicate significantly elevated CaMK, increased SERCA activity and significantly reduced I_to_ as the key contributors to HCM pathophysiology. These combine to elevate the [𝐶𝑎^2+^]_𝑖_ and lead to increased ATP consumption to maintain contractility. CaMK, SERCA and I_to_ thus serve as potential therapeutic targets and diagnostic or prognostic markers for HCM.

## Data Availability

The data and source code are available by contacting the corresponding author upon request.

## Acknowledgments

This manuscript has also been presented in bioRxiv (Adeniran et al. 2023).

## Supplementary Materials

Figure showing the crossbridge states resulting from a second [𝐶𝑎^2+^]_𝑖_ transient profile from HCM patients with a MYBPC3 mutation using the spatial multifilament, half sarcomere model.

## Supplementary Materials

**Figure S1:**
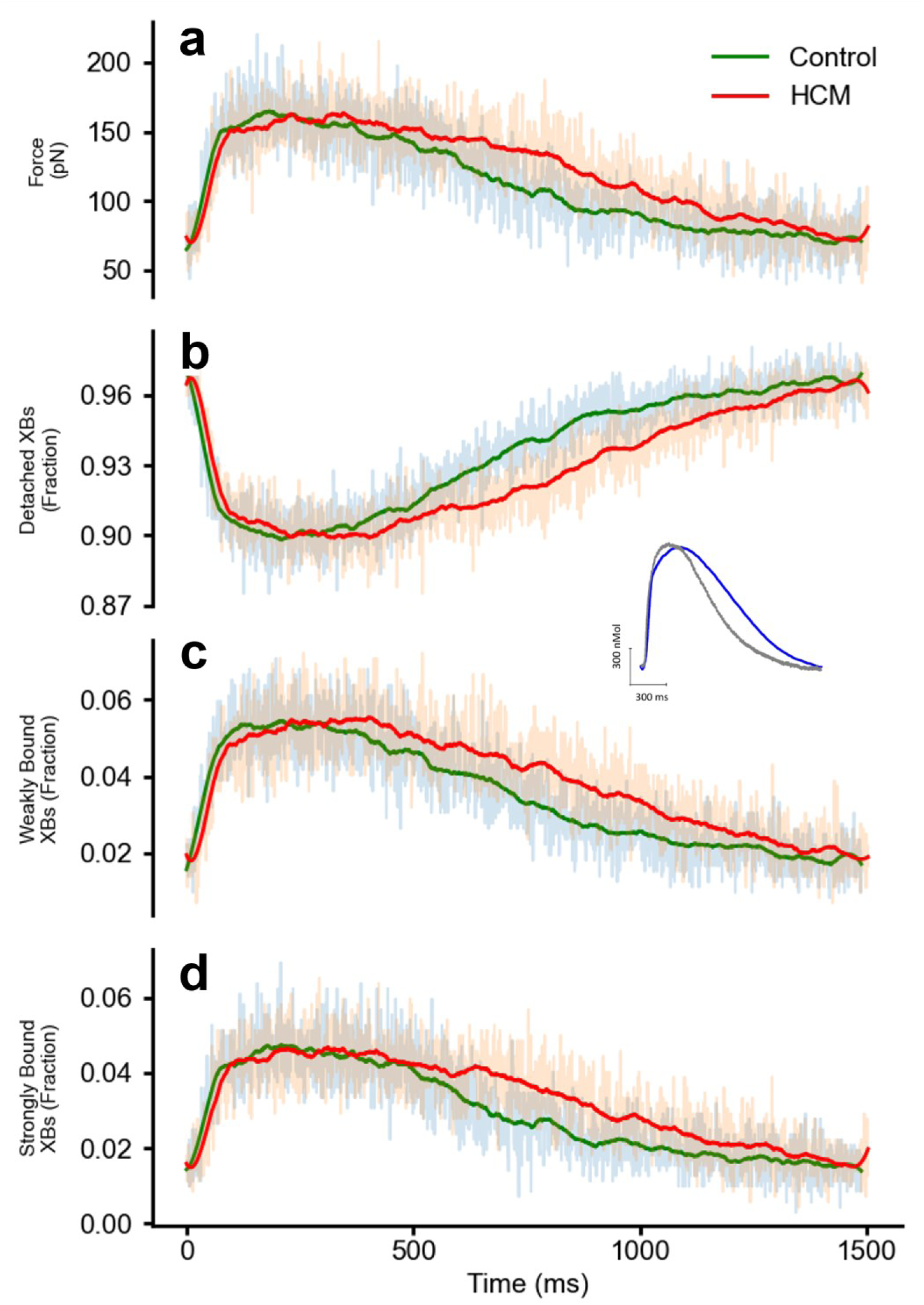
Crossbridge states in control and HCM (MYBPC3 mutations) with the spatial multifilament, half sarcomere model. (a) The developed force in control (green) and HCM (red). (b) The fraction of crossbridges in the detached state during the time course of the [𝐶𝑎^2+^]_𝑖_ transient profile. (c) The fraction of crossbridges in the weakly bound state during the time course of the [𝐶𝑎^2+^]_𝑖_ transient profile. (d) The fraction of crossbridges in the strongly bound state during the time course of the [𝐶𝑎^2+^]_𝑖_ transient profile.

## References

1. Ashrafian, H., Redwood, C., Blair, E., Watkins, H., 2003. Hypertrophic cardiomyopathy:a paradigm for myocardial energy depletion. Trends Genet. 19, 263–268. 10.1016/S0168-9525(03)00081-7

2. Chung, J.-H., Biesiadecki, B.J., Ziolo, M.T., Davis, J.P., Janssen, P.M.L., 2016. Myofilament Calcium Sensitivity: Role in Regulation of In vivo Cardiac Contraction and Relaxation. Front. Physiol. 7, 562. 10.3389/fphys.2016.00562

3. Coppini, R., Ferrantini, C., Mugelli, A., Poggesi, C., Cerbai, E., 2018. Altered Ca2+ and Na+ Homeostasis in Human Hypertrophic Cardiomyopathy: Implications for Arrhythmogenesis. Front. Physiol. 9. 10.3389/fphys.2018.01391

4. Coppini, R., Ferrantini, C., Yao, L., Fan, P., Del Lungo, M., Stillitano, F., Sartiani, L., Tosi, B., Suffredini, S., Tesi, C., Yacoub, M., Olivotto, I., Belardinelli, L., Poggesi, C., Cerbai, E., Mugelli, A., 2013. Late Sodium Current Inhibition Reverses Electromechanical Dysfunction in Human Hypertrophic Cardiomyopathy. Circulation 127, 575–584. 10.1161/CIRCULATIONAHA.112.134932

5. Cortassa, S., Aon, M.A., Marbán, E., Winslow, R.L., O’Rourke, B., 2003. An integrated model of cardiac mitochondrial energy metabolism and calcium dynamics. Biophys. J. 84, 2734–2755. 10.1016/s0006-3495(03)75079-6

6. Del Franco, A., Ambrosio, G., Baroncelli, L., Pizzorusso, T., Barison, A., Olivotto, I., Recchia, F.A., Lombardi, C.M., Metra, M., Ferrari Chen, Y.F., Passino, C., Emdin, M., Vergaro, G., 2022. Creatine deficiency and heart failure. Heart Fail. Rev. 27, 1605–1616. 10.1007/s10741-021-10173-y

7. Forouzandehmehr M, Paci M, Koivumäki JT, Hyttinen J. 2022. Altered contractility in mutation-specific hypertrophic cardiomyopathy: A mechano-energetic in silico study with pharmacological insights. Front Physiol. 13:1010786. 10.3389/fphys.2022.1010786.

8. Haland, T.F., Hasselberg, N.E., Almaas, V.M., Dejgaard, L.A., Saberniak, J., Leren, I.S., Berge, K.E., Haugaa, K.H., Edvardsen, T., 2017. The systolic paradox in hypertrophic cardiomyopathy. Open Heart 4, e000571. 10.1136/openhrt-2016-000571

9. Helms, A.S., Tang, V.T., O’Leary, T.S., Friedline, S., Wauchope, M., Arora, A., Wasserman, A.H., Smith, E.D., Lee, L.M., Wen, X.W., Shavit, J.A., Liu, A.P., Previs, M.J., Day, S.M., 2020. Effects of MYBPC3 loss-of-function mutations preceding hypertrophic cardiomyopathy. JCI Insight 5. 10.1172/jci.insight.133782

10. Henry, J.A., Levelt, E., Rayner, J., Hundertmark, M., Peterzan, M., Green, P., Watson, W., Lewis, A., Burrage, M., Arvidsson, P., Chamley, R., Nicol, E., Holdsworth, D., Neubauer, S., Valkovic, L., Rider, O., 2022. 143 Measuring pcr/atp as a marker of myocardial energetics across the spectrum of metabolic cardiac disease. Heart 108, A109–A110. 10.1136/heartjnl-2022-BCS.143

11. Hindmarsh, A., Petzold, L.R., 2005. 143 LSODA, Ordinary Differential Equation Solver for Stiff or Non-Stiff System. (Sep 2005). Bibliographic information available from INIS: http://inis.iaea.org/search/search.aspx?orig_q=RN:41086668 Available on-line: http://www.nea.fr/abs/html/uscd1227.html

12. Li, S., Pan, H., Tan, C., Sun, Y., Song, Y., Zhang, X., Yang, W., Wang, X., Li, D., Dai, Y., Ma, Q., Xu, C., Zhu, X., Kang, L., Fu, Y., Xu, X., Shu, J., Zhou, N., Han, F., Qin, D., Huang, W., Liu, Z., Yan, Q., 2018. Mitochondrial Dysfunctions Contribute to Hypertrophic Cardiomyopathy in Patient iPSC-Derived Cardiomyocytes with MT-RNR2 Mutation. Stem Cell Rep. 10, 808–821. 10.1016/j.stemcr.2018.01.013

13. Liu T, Li X, Wang Y, Zhou M, Liang F. 2023. Computational modeling of electromechanical coupling in human cardiomyocyte applied to study hypertrophic cardiomyopathy and its drug response. Comput Methods Programs Biomed. 231:107372. 10.1016/j.cmpb.2023.107372.

14. Lucas, D.T., Aryal, P., Szweda, L.I., Koch, W.J., Leinwand, L.A., 2003. Alterations in mitochondrial function in a mouse model of hypertrophic cardiomyopathy. Am. J. Physiol.-Heart Circ. Physiol. 284, H575–H583. 10.1152/ajpheart.00619.2002

15. Lymn, R.W., Taylor, E.W., 1971. Mechanism of adenosine triphosphate hydrolysis by actomyosin. Biochemistry 10, 4617–4624. 10.1021/bi00801a004

16. Marian, A.J., Braunwald, E., 2017. Hypertrophic Cardiomyopathy: Genetics, Pathogenesis, Clinical Manifestations, Diagnosis, and Therapy. Circ. Res. 121, 749–770. 10.1161/CIRCRESAHA.117.311059

17. Neubauer S, Horn M, Cramer M et al., 1997. Myocardial phosphocreatine-to-ATP ratio is a predictor of mortality in patients with dilated cardiomyopathy. Circulation 96, 2190–2196.

18. O’Hara, T., Virág, L., Varró, A., Rudy, Y., 2011. Simulation of the Undiseased Human Cardiac Ventricular Action Potential: Model Formulation and Experimental Validation. PLOS Comput. Biol. 7, e1002061. 10.1371/journal.pcbi.1002061

19. Ormerod, J.O.M., Frenneaux, M.P., Sherrid, M.V., 2016. Myocardial energy depletion and dynamic systolic dysfunction in hypertrophic cardiomyopathy. Nat. Rev. Cardiol. 13, 677–687. 10.1038/nrcardio.2016.98

20. Pasipoularides, A., 2018. Challenges and Controversies in Hypertrophic Cardiomyopathy: Clinical, Genomic and Basic Science Perspectives. Rev. Espanola Cardiol. Engl. Ed 71, 132–138. 10.1016/j.rec.2017.07.003

21. Pioner, J.M., Vitale, G., Steczina, S., Langione, M., Margara, F., Santini, L., Giardini, F., Lazzeri, E., Piroddi, N., Scellini, B., Palandri, C., Schuldt, M., Spinelli, V., Girolami, F., Mazzarotto, F., van der Velden, J., Cerbai, E., Tesi, C., Olivotto, I., Bueno-Orovio, A., Sacconi, L., Coppini, R., Ferrantini, C., Regnier, M., Poggesi, C., 2023. Slower Calcium Handling Balances Faster Cross-Bridge Cycling in Human MYBPC3 HCM. Circ. Res. 132, 628–644. 10.1161/CIRCRESAHA.122.321956

22. Powers, J.D., Williams, C.D., Regnier, M., Daniel, T.L., 2018. A Spatially Explicit Model Shows How Titin Stiffness Modulates Muscle Mechanics and Energetics. Integr. Comp. Biol. 58, 186–193. 10.1093/icb/icy055

23. Ranjbarvaziri, S., Kooiker, K.B., Ellenberger, M., Fajardo, G., Zhao, M., Vander Roest, A.S., Woldeyes, R.A., Koyano, T.T., Fong, R., Ma, N., Tian, L., Traber, G.M., Chan, F., Perrino, J., Reddy, S., Chiu, W., Wu, J.C., Woo, J.Y., Ruppel, K.M., Spudich, J.A., Snyder, M.P., Contrepois, K., Bernstein, D., 2021. Altered Cardiac Energetics and Mitochondrial Dysfunction in Hypertrophic Cardiomyopathy. Circulation 144, 1714–1731. 10.1161/CIRCULATIONAHA.121.053575

24. Rice, J.J., Wang, F., Bers, D.M., de Tombe, P.P., 2008. Approximate Model of Cooperative Activation and Crossbridge Cycling in Cardiac Muscle Using Ordinary Differential Equations. Biophys. J. 95, 2368– 2390. 10.1529/biophysj.107.119487

25. Rowin, E.J., Fifer, M.A., 2021. Evaluating Histopathology to Improve Our Understanding of Hypertrophic Cardiomyopathy∗. J. Am. Coll. Cardiol. 77, 2171–2173. 10.1016/j.jacc.2021.03.292

26. Santo-Domingo, J., Demaurex, N., 2010. Calcium uptake mechanisms of mitochondria. Biochim. Biophys. Acta BBA - Bioenerg., 16th European Bioenergetics Conference 2010 1797, 907–912. 10.1016/j.bbabio.2010.01.005

27. Semsarian, C., 2018. Update on the Diagnosis and Management of Hypertrophic Cardiomyopathy. Heart Lung Circ. 27, 276–279. 10.1016/j.hlc.2017.10.007

28. Spudich, J.A., 2014. Hypertrophic and Dilated Cardiomyopathy: Four Decades of Basic Research on Muscle Lead to Potential Therapeutic Approaches to These Devastating Genetic Diseases. Biophys. J. 106, 1236– 1249. 10.1016/j.bpj.2014.02.011

29. Sun, Y.-B., Irving, M., 2010. The molecular basis of the steep force–calcium relation in heart muscle. J. Mol. Cell. Cardiol. 48, 859–865. 10.1016/j.yjmcc.2009.11.019

30. Tardiff, J.C., Hewett, T.E., Palmer, B.M., Olsson, C., Factor, S.M., Moore, R.L., Robbins, J., Leinwand, L.A., 1999. Cardiac troponin T mutations result in allele-specific phenotypes in a mouse model for hypertrophic cardiomyopathy. J. Clin. Invest. 104, 469–481. 10.1172/JCI6067

31. Tejado, B.S.M., Jou, C., n.d. Histopathology in HCM. Glob. Cardiol. Sci. Pract. 2018, 20. 10.21542/gcsp.2018.20

32. Tran, K., Smith, N.P., Loiselle, D.S., Crampin, E.J., 2010. A Metabolite-Sensitive, Thermodynamically Constrained Model of Cardiac Cross-Bridge Cycling: Implications for Force Development during Ischemia. Biophys. J. 98, 267–276. 10.1016/j.bpj.2009.10.011

33. Unno, K., Isobe, S., Izawa, H., Cheng, X.W., Kobayashi, M., Hirashiki, A., Yamada, T., Harada, K., Ohshima, S., Noda, A., Nagata, K., Kato, K., Yokota, M., Murohara, T., 2009. Relation of functional and morphological changes in mitochondria to myocardial contractile and relaxation reserves in asymptomatic to mildly symptomatic patients with hypertrophic cardiomyopathy. Eur. Heart J. 30, 1853– 1862. 10.1093/eurheartj/ehp184

34. Wijnker, P.J.M., Sequeira, V., Kuster, D.W.D., van der Velden, J., 2019. Hypertrophic Cardiomyopathy: A Vicious Cycle Triggered by Sarcomere Mutations and Secondary Disease Hits. Antioxid. Redox Signal. 31, 318–358. 10.1089/ars.2017.7236

35. Williams, C.D., Regnier, M., Daniel, T.L., 2012. Elastic Energy Storage and Radial Forces in the Myofilament Lattice Depend on Sarcomere Length. PLOS Comput. Biol. 8, e1002770. 10.1371/journal.pcbi.1002770

36. Williams, C.D., Regnier, M., Daniel, T.L., 2010. Axial and Radial Forces of Cross-Bridges Depend on Lattice Spacing. PLOS Comput. Biol. 6, e1001018. 10.1371/journal.pcbi.1001018

37. Williams, C.D., Salcedo, M.K., Irving, T.C., Regnier, M., Daniel, T.L., 2013. The length–tension curve in muscle depends on lattice spacing. Proc. R. Soc. B Biol. Sci. 280, 20130697. 10.1098/rspb.2013.0697

38. Zorova, L.D., Popkov, V.A., Plotnikov, E.Y., Silachev, D.N., Pevzner, I.B., Jankauskas, S.S., Babenko, V.A., Zorov, S.D., Balakireva, A.V., Juhaszova, M., Sollott, S.J., Zorov, D.B., 2018. Mitochondrial membrane potential. Anal. Biochem. 552, 50–59. 10.1016/j.ab.2017.07.009

